# NTxPred2: A large language model for predicting neurotoxic peptides and neurotoxins

**DOI:** 10.1101/2025.03.01.640936

**Authors:** Anand Singh Rathore, Saloni Jain, Shubham Choudhury, Gajendra P. S. Raghava

**Author notes:** **Mailing Address of Authors** Anand Singh Rathore (ASR), Saloni Jain (SJ), Shubham Choudhury (SC), Gajendra P. S. Raghava (GPSR). **Corresponding Author** Prof. Gajendra P. S. Raghava Head and Professor, Department of Computational Biology, Indraprastha Institute of Information Technology, Delhi, Okhla Industrial Estate, Phase III, (Near Govind Puri Metro Station) New Delhi, India – 110020, Office: A-302 (R&D Block), Website: http://webs.iiitd.edu.in/raghava/. **Author’s Biography** 1. Anand Singh Rathore is currently pursuing a Ph.D. in Computational Biology at the Department of Computational Biology, Indraprastha Institute of Information Technology, New Delhi, India. 2. Salini Jain is currently pursuing a Ph.D. in Computational Biology at the Department of Computational Biology, Indraprastha Institute of Information Technology, New Delhi, India. 3. Shubham Choudhury is currently pursuing a Ph.D. in Computational Biology at the Department of Computational Biology, Indraprastha Institute of Information Technology, New Delhi, India.

## Abstract

Most of the existing methods including our NTxPred, use a single model to predict both neurotoxic peptides and neurotoxic proteins (neurotoxins). In this study, we developed distinct models for predicting neurotoxic peptides and neurotoxins. Our peptide dataset consists of 877 neurotoxic and 877 non-neurotoxic peptides, while our protein dataset includes 775 neurotoxins and 775 non-neurotoxins. Preliminary analysis reveals that certain residues, such as cysteine, are more prevalent in both neurotoxic peptides and proteins, though their abundance differs in magnitude. First, we developed machine learning models using composition and binary profiles, achieving a maximum AUC of 0.97 for peptides and 0.85 for proteins. The performance for proteins improved from an AUC of 0.85 to 0.89 when evolutionary information was incorporated. Next, we built machine learning models using embeddings from protein language models, attaining an AUC of 0.96 for peptides and 0.94 for proteins. We also developed protein language models and achieved an AUC of 0.98 for peptides using esm2-t30 and 0.91 for proteins using esm2-t6. All models were trained and tested using 5-fold cross-validation, and final models were evaluated on an independent dataset not used in training. We further assessed protein models on the peptide dataset and vice versa, highlighting the necessity of separate models. The proposed models outperform existing methods on independent datasets. Our neurotoxicity prediction models will aid in the safety assessment of genetically modified foods and therapeutic proteins by minimizing the need for animal testing. To support the scientific community, we developed a standalone software and web server NTxPred2, for predicting and scanning neurotoxins (https://webs.iiitd.edu.in/raghava/ntxpred2/, https://github.com/raghavagps/ntxpred2/).

**Highlights:**

- Development of models for predicting neurotoxic peptides and neurotoxins
- ML Models Using LLM Embeddings for Neurotoxin Prediction
- Protein language models esm2-t30 for predicting neurotoxic peptides
- Important for risk assessment of genetically modified foods/proteins
- NTxPred2 is an updated version of NTxPred for predicting neurotoxicity

## 1 Introduction

Over the past 150 years, significant advancements have been made in utilizing proteins for therapeutic applications. A major breakthrough occurred in 1891 with the development of an antibody-based treatment for diphtheria, which led to the awarding of the first Nobel Prize in Medicine [1]. This was followed by the approval of insulin, a peptide-based drug, in 1922, providing a life-saving treatment for diabetes. Initially, insulin production relied on animal-derived sources due to limitations in human insulin production. However, the advent of recombinant DNA technology revolutionized this process. In 1982, insulin was successfully produced in E. coli using recombinant techniques, making it the first FDA-approved recombinant protein therapy [2]. Today, peptide and protein-based drugs constitute over 10% of the pharmaceutical market, with their share expected to grow in the coming years. The Therapeutic Peptide and Protein Database (THPdb) serves as a comprehensive resource for US FDA-approved protein and peptide therapeutics [3]. The first version, published in 2017, documented 239 approved therapeutic peptides and proteins. The latest version, THPdb2, released in 2024, has expanded to include 894 FDA-approved protein-based therapeutics, highlighting the importance of peptides and proteins in modern medicine [4]. Additionally, genetically engineered organisms and crops are being used to produce edible proteins at a large scale. The World Health Organization (WHO) has established guidelines for evaluating the safety of both edible proteins and protein-based therapeutics. One of the major challenges in these guidelines is assessing the potential adverse effects, particularly toxicities, of newly discovered proteins. Though a number of experimental techniques have been developed to assess protein-related toxicities, these methods are expensive, time-consuming, and labor-intensive. To support experimental researchers, numerous in silico tools have been developed to screen proteins that have any type of toxicity like cytotoxicity, immunotoxicity, hemotoxicity, and neurotoxicity [5–9].

In the last one-decade number of methods have been developed for predicting the cytotoxicity of peptides and proteins it including ToxinPred [9–11], ToxClassifier [12], TOXIFY[13], ToxDL [14], ATSE [15], ToxIBTL [16], ToxMVA [17], CSM toxin [18], VISHpred [19], and MultiToxPred 1.0 [20]. Similarly, a number of in silico tools have been developed for predicting the allergenicity or immunotoxicity of proteins include IL6pred [7] and AlgPred [21]. Despite these advancements, limited efforts have been made to predict the neurotoxicity of proteins and peptides. Neurotoxicity is a type of toxicity that causes damage to the structure and function of the brain, spinal cord, and peripheral nervous systems. It has been a number of studies that a number of proteins may also have neurotoxicity, for example, diphtheria toxin, apamin, palytoxin, and scorpion toxins [22–26]. Thus, neurotoxicity is a significant concern in the field of protein therapeutics and in genetically modified foods. In order to address this issue, Saha and Raghava developed a method NTxPred in year 2017, for identifying neurotoxins based on their functional and source characteristics [27]. This tool has found applications in various forms in the development of therapeutic peptides [28,29]. Guang et al. (2010) developed an SVM-based method using evolutionary information for predicting neurotoxins [30]. In addition, numerous methods have been developed for the prediction and characterization of spider neurotoxins [31,32]. Neurotoxins are categorized into presynaptic and postsynaptic neurotoxins based on their synaptic location and mechanism of action. A range of machine-learning techniques has been developed to classify presynaptic and postsynaptic neurotoxins [33–40]. In Supplementary Table S1, we provide a comprehensive overview of methodologies employed for predicting specific toxicity types in therapeutic peptides and proteins, including hemotoxicity [41–48].

This study aims to establish a highly precise and dependable approach for predicting the toxicity of peptides and proteins. Our previous method, NTxPred, introduced in 2007, was developed using a limited dataset and encountered challenges due to the scarce availability of neurotoxic peptide and protein data. Consequently, it offered a single predictive model for both categories, without specialized strategies tailored to peptides and proteins individually. Over the past two decades, researchers have annotated numerous neurotoxic peptides and proteins, which have been deposited in Swiss-Prot database. To address the limitations of our previous study, we extracted experimentally validated neurotoxic peptides and proteins from the latest release of the Swiss-Prot. This new dataset contains several times more proteins and peptides compared to the dataset used in NTxPred. Consequently, we made a systematic effort to develop distinct protein-specific and peptide-specific models for neurotoxicity prediction. Additionally, we conducted cross-dataset analysis to assess whether a unified model trained on both peptides and proteins could outperform models trained separately on individual datasets. To achieve this, we used the largest available experimentally validated dataset and incorporated cutting-edge algorithmic frameworks, including SOTA transformer-based language models and ensemble learning techniques [61]. These advancements enhance the robustness of the model, ensure comprehensive feature representation, and significantly improve predictive accuracy. By integrating these innovations, our upgraded NTxPred2 method aims to provide a highly reliable tool for neurotoxicity prediction, contributing to the safer design and development of protein/peptide-based drugs.

## 2 Methods

### 2.1 Data Collection and Pre-processing

A comprehensive dataset of experimentally validated neurotoxic and non-toxic peptides was curated from the Swiss-Prot. To construct the positive dataset, the keyword “neurotoxin” was queried with the “reviewed_true” filter, retrieving an initial set of 5173 putative neurotoxic peptide sequences. For the negative dataset, sequences were obtained using the keyword “NOT toxin NOT neurotoxin” with the “reviewed_true” filter, yielding an initial set of 566303 non-toxic peptide sequences. This information has been used to create the following three datasets:

1. Peptide Dataset: In this case, we only select peptides having ≤ 51 amino acids. In addition, we removed peptides containing non-canonical amino acids and redundant peptides. Finally, we got 877 unique neurotoxic peptides called positive dataset of peptides. The same exclusion criteria were applied to non-neurotoxic peptides, yielding 8,437 unique non-toxic peptides. To mitigate the class imbalance inherent in the dataset, the CD-HIT software [62] was employed to cluster the non-toxic peptide sequences at a 60% sequence identity threshold. This clustering significantly reduced the number of redundant sequences within the negative dataset. Resulting in a negative dataset of 877 non-neurotoxic peptides with length distributions matching the positive dataset. Although CD-HIT 40 was also considered, it did not yield a sufficient number of non-toxic peptides to maintain length balance. The minimum peptide length included in the dataset was 7 amino acids.
2. Protein Dataset: In case of protein dataset, we only select proteins having ≥ 51 amino acids. The same filtering criteria were applied, removing sequences with non-canonical amino acids, duplicates, and redundant entries, resulting in 3,755 neurotoxic and 457745 non-toxic protein sequences. To further refine the dataset and reduce redundancy, CD-HIT 40 was applied, yielding 775 non-redundant neurotoxic protein sequences. To create a balanced dataset, 775 non-toxic protein sequences of comparable lengths were selected.

To ensure a rigorous and unbiased model evaluation, standard data partitioning strategies were employed. Both the peptide and protein datasets were independently divided into two subsets. The training set (80% of the data) is used for model development, and the independent dataset (20% of the data) is reserved strictly for final model evaluation. It is important that the independent dataset remains completely unseen during training, testing, model selection, and hyperparameter tuning to prevent data leakage and ensure an accurate assessment of model generalizability.

3. Combined Dataset: To evaluate the model’s ability to generalize across peptide and protein sequences, a combined dataset was created. Instead of simply merging all peptide and protein sequences, this dataset was structured such that the training datasets of peptides and proteins were merged to form a combined training dataset. The independent datasets of peptides and proteins were merged to form a combined independent dataset for model evaluation. This approach ensures a fair and robust model assessment by evaluating performance on a truly independent dataset containing both molecular types.

The complete data curation process is illustrated in Figure 1.

**Figure 1:**
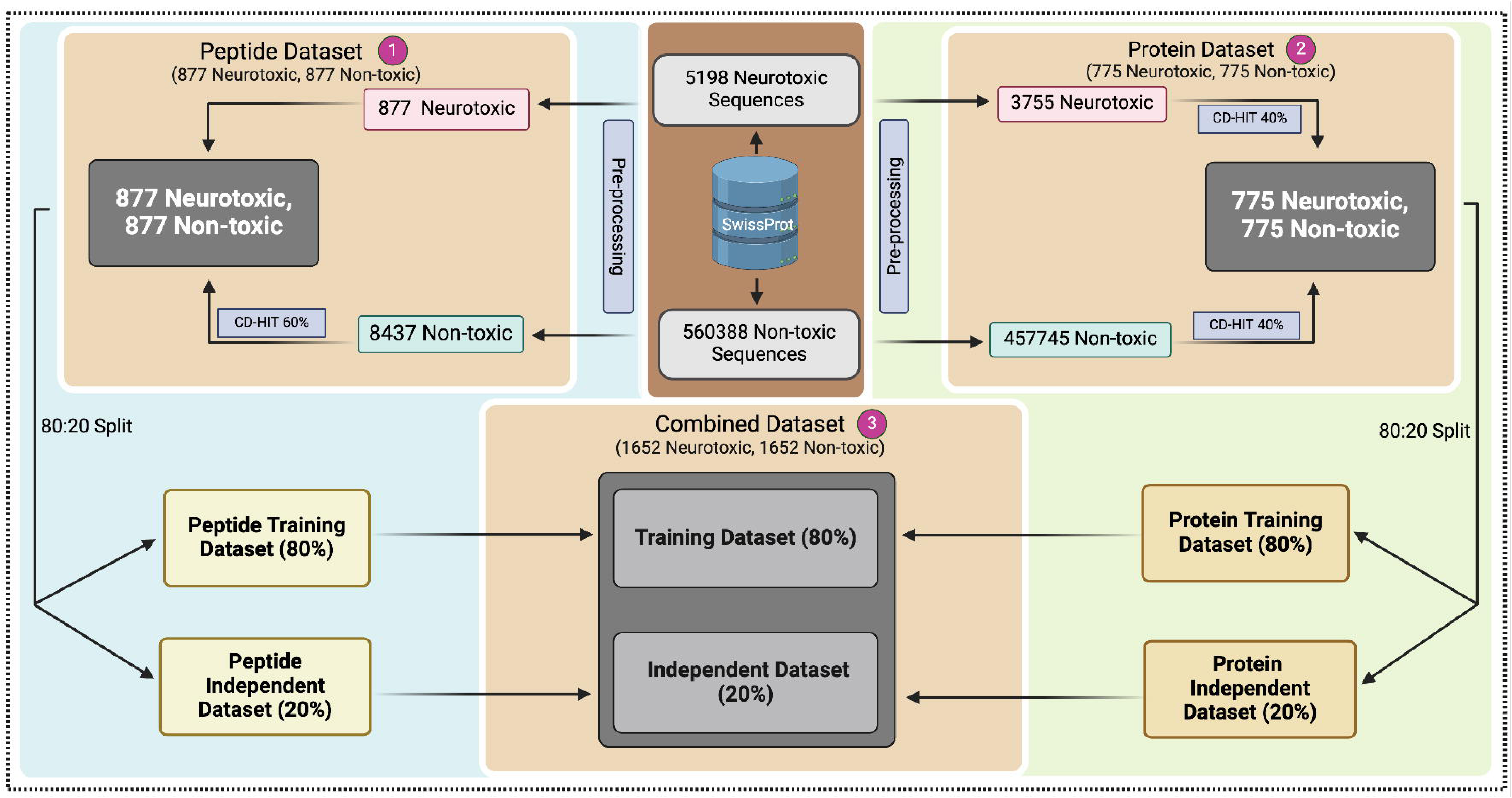
A schematic representation of NTXPred2 dataset creation.

### 2.2 Compositional and Positional Analysis

To gain an initial understanding of the distinguishing features of experimentally validated neurotoxic and non-toxic sequences, we conducted a comprehensive compositional and positional analysis. Our approach involved multiple analytical techniques to assess sequence-based properties across all three datasets. The first stage of analysis focused on amino acid composition within each class. For both neurotoxic and non-toxic sequences, we calculated key statistical metrics, including the mean, median, and standard deviation of amino acid frequencies. To assess the statistical significance of differences between the two classes, we employed independent t-tests (*scipy.stats*) and controlled for false discovery rates using the Benjamini-Hochberg procedure (*statsmodels.stats.multitest*). These statistical corrections ensured robustness by minimizing type I errors. Beyond the neurotoxic and non-toxic classifications, we extended our analysis to compare amino acid compositions across other toxicity categories, including hemotoxicity [48] and cytotoxicity [9], as well as a general genome-wide amino acid composition. This broader comparison provided insights into the shared and unique sequence characteristics of different toxicity classes. To further investigate sequence-specific preferences, we applied the Two Sample Logo (TSL) method, enabling the identification of distinct positional preferences for amino acid residues. This technique highlighted residues that were preferentially enriched or depleted at specific positions within peptide sequences, offering deeper insights into sequence associated with neurotoxicity.

### 2.3 Correlation Between Toxicities

To explore the relationships between different toxicity types, we conducted a correlation analysis based on amino acid composition patterns. This analysis aimed to identify potential overlaps or distinctions in compositional features among various toxicity classes, including neurotoxicity, hemotoxicity, and cytotoxicity. By systematically comparing the amino acid composition profiles of these toxicity categories, we sought to determine whether certain residues were consistently associated with toxicity across multiple classes or if unique compositional signatures distinguished one toxicity type from another. This investigation provides valuable insights into the underlying molecular characteristics that drive toxicity and helps in understanding the extent to which different toxic properties share common sequence determinants.

### 2.4 Composition and Physicochemical Property-based Prediction

In this analysis, we classified neurotoxic and non-toxic sequences based on the composition of each of the 20 standard amino acids and different types of physicochemical properties like polarity, charge, etc. For each feature, we first calculated the mean composition for both the neurotoxic and non-toxic groups. A threshold was then determined as the midpoint between the two means. Sequences were classified according to whether their composition exceeded this threshold. Specifically, if the neurotoxic group had a higher mean composition for a particular amino acid, sequences with values above the threshold were classified as neurotoxic, and vice versa. This classification was performed across all three datasets. To assess the classification performance, we calculated the accuracy and area under the curve (AUC) for each amino acid. The results, including the thresholds, mean compositions, accuracy, and AUC, were compiled into a comprehensive table for further analysis.

### 2.4 Features Extraction

In developing a sequence-based predictor for peptide and protein properties, it is crucial to effectively represent sequences to capture their biological functions. We extracted a variety of informative features, including binary profiling, PSSM profiling, composition-based features, physiochemical features, and word embeddings using large language models (LLMs). These features were computed across all three datasets to improve the predictive accuracy and generalizability of the model.

#### 2.4.1 Binary Profiling

To assess the importance of specific residues at the N- and C-terminal positions, we incorporated both frequency and positional information of residues. We developed a method using binary profiles of peptides, generating these profiles for each sequence across all three datasets. Since the minimum sequence length was seven, we extracted the first seven amino acids (NT7) and the last seven amino acids (CT7) from each sequence. Each amino acid in these regions was then one-hot encoded, creating a 14-dimensional binary vector that indicated the presence or absence of the 20 standard amino acids at each position. As a result, each sequence was represented by a 280-dimensional feature vector, capturing the amino acid composition of its terminal regions. These binary profiles were integrated into the dataset, providing valuable features for subsequent machine-learning analyses. This approach has been successfully applied in several existing methods [63,64].

#### 2.4.2 Evolutionary Information-based Features

Evolutionary information-based features capture sequence conservation and variability, providing insights into functional importance [65,66]. In this study, we utilized the Position-Specific Scoring Matrix (PSSM), obtained through Position-Specific Iterated BLAST (PSI-BLAST) [67], to extract evolutionary information. The PSSM matrix quantifies the evolutionary conservation of each amino acid position in a sequence by comparing it to a set of homologous sequences. We employed multiple modules from the *POSSUM* package [68]:

1. *aac_pssm*: This module calculates the average activation values for each amino acid position across the sequence, providing a 20-dimensional (20D) feature vector. It captures how strongly each amino acid type is represented at each position, reflecting sequence-specific characteristics.
2. *pssm_composition*: This method treats the entire PSSM matrix as a single vector and computes its average, encoding global properties of the matrix. It results in a 400-dimensional (400D) feature vector that reflects the overall evolutionary profile of the sequence.
3. *mepd_pssm* (Mean Evolutionary Difference Profile): This combines features from two approaches-Evolutionary Difference Profile (EDP) and Evolutionary Difference PSSM (EEDP). EDP computes differences between PSSM values for paired residues across rows and columns, highlighting evolutionary variation, while EEDP focuses on evolutionary differences between PSSM values for paired positions, distinguishing conserved from variable regions. The *mepd_pssm* method yields a 420-dimensional (420D) feature vector.

For this study, we generated PSSM matrices for each sequence using Swiss-Prot as the reference database. However, two sequences were excluded from the analysis due to the absence of homologous sequences, preventing PSSM generation. Additionally, we applied these features only to the protein dataset, as PSSM-based methods are generally unreliable for short sequences. Specifically, sequences shorter than 50 residues lack sufficient evolutionary context, leading to poorly constructed PSSM matrices with high variability and low statistical confidence [69–71].

#### 2.4.3 Composition-based Features

For composition-based feature extraction, we used the standalone tool of *Pfeature*, which provided a wide array of descriptors. In total, we collected 9,190 features for each peptide, encompassing 18 distinct feature types. These types include Amino Acid Composition (AAC), Dipeptide Composition (DPC), Atom Type Composition, Bond Type Composition, and various physicochemical properties (PCP), among others. Detailed descriptions of each feature and the corresponding vector lengths are presented in Supplementary Table S2.

In addition to Pfeature, we also utilized the modlAMP library. We calculated 56 physicochemical properties for each sequence based on their primary sequences, utilizing the *modlAMP* 4.3.0 package in Python [72]. This included 47 sequence-specific descriptors and 9 global descriptors. A comprehensive list of these properties can be found in Supplementary Table S3. The resulting data was organized into datasets with peptide sequences as rows and their corresponding properties as columns. This approach has been employed in various studies focused on toxicity prediction [73].

#### 2.4.4 Word Embeddings

Recent advancements in natural language processing (NLP) have led to the development of Protein Language Models (PLMs), which utilize amino acids and their combinations (doublets or triplets) as tokens. These models generate fixed-size vectors, known as embeddings, which encapsulate specific peptide sequences. These protein embeddings are crucial for various applications, including structure prediction, novel sequence generation, and protein classification [74,75]. In our research, we utilized two prominent large language models (LLMs): esm-2 [76] and ProtBERT. These PLMs are recognized as a state-of-the-art model for predicting various protein properties directly from individual sequences [19,77]. ProtBERT is an extension of the BERT model, pre-trained on extensive protein sequence datasets using a self-supervised approach [78]. esm-2, a transformer-based PLM, was trained on sequences from the UniRef protein sequence database using a masked language modeling objective. Figure 2 showcases different protein language models used to extract the embeddings, each varying in size and complexity. Several checkpoints of the esm-2 model, which vary in size, can be found on Hugging Face [79]. Generally, larger models tend to achieve marginally improved accuracy; however, they also demand considerably more memory and longer training durations. Key characteristics such as the number of layers, the total number of parameters in which they are trained, and the embedding dimension are displayed for each model. This visualization provides a concise overview of these protein language models, highlighting their key specifications.

**Figure 2:**
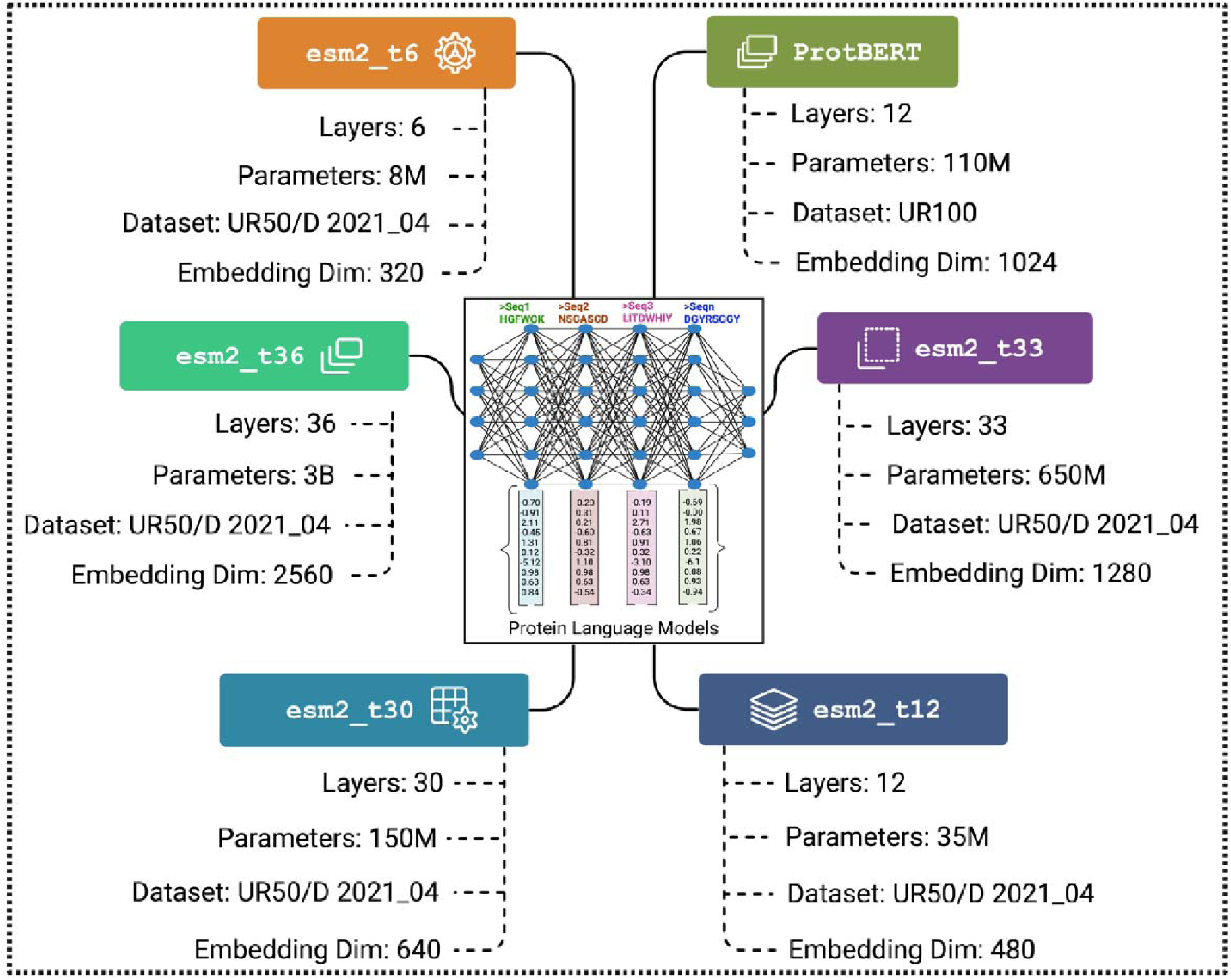
Protein Language Models: A Visual Overview.

### 2.5 Machine Learning-based classifiers

Machine learning algorithms have been extensively utilized to distinguish neurotoxic peptides and proteins from non-toxic ones. In this study, we explored a range of classifiers, including Logistic Regression (LR), Ridge, Lasso, ElasticNet, Random Forest (RF), Extra Trees (ET), Gaussian Naïve Bayes (GNB), Decision Trees (DT), Multi-Layer Perceptron (MLP), XGBoost (XGB), AdaBoost (AdB), and Support Vector Classifier (SVC) with various kernel functions (linear, RBF, polynomial, and sigmoid). Each model was systematically optimized by fine-tuning multiple hyperparameters to achieve optimal classification performance. The complete workflow of the NTxPred2 system is depicted in Figure 3.

**Figure 3:**
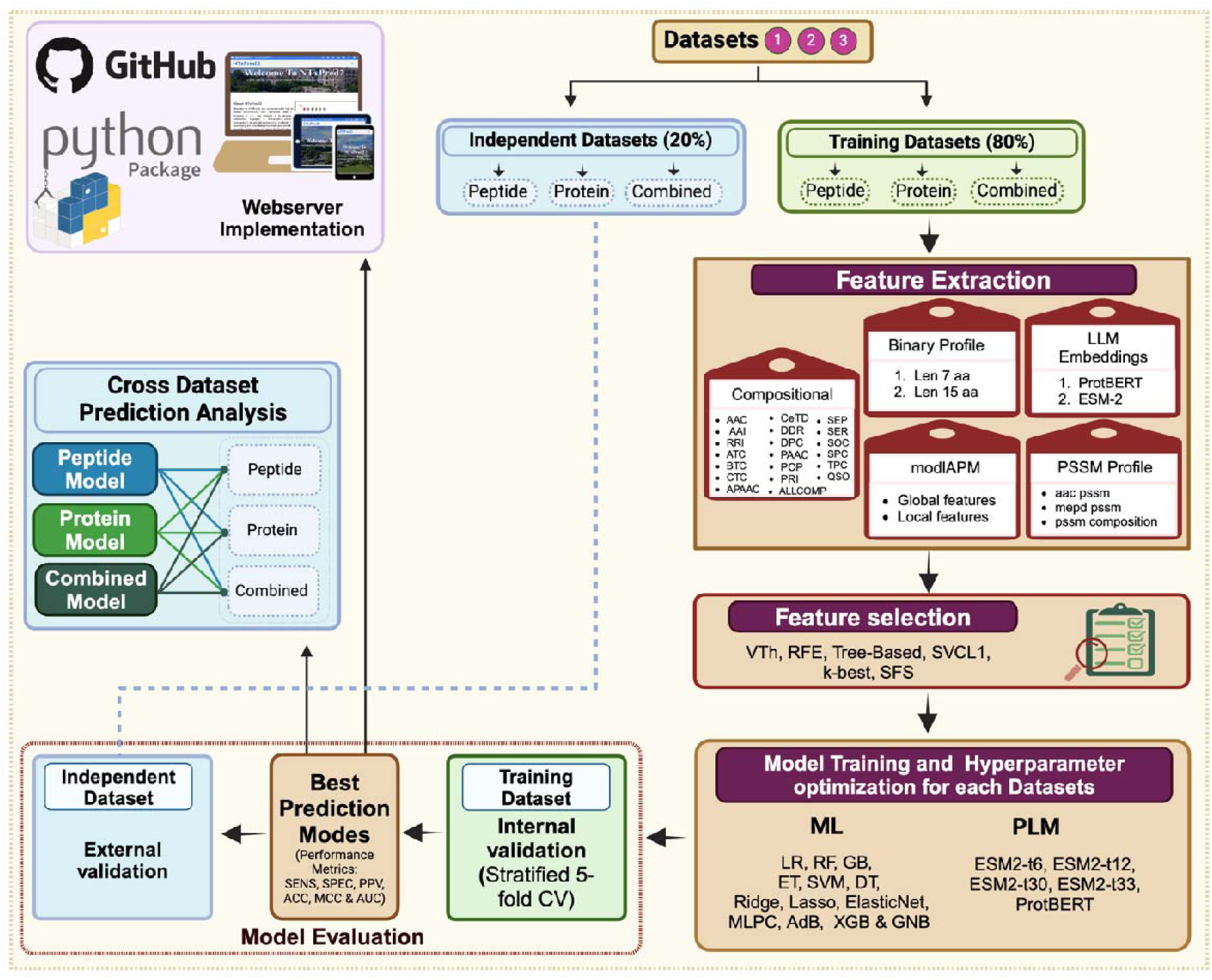
Flowchart illustrating the complete architecture of NTxPred2.

### 2.6 Feature Selection

Feature selection is a crucial step in machine learning that enhances model performance by identifying and retaining only the most informative features while eliminating redundant or irrelevant ones. This process not only improves prediction accuracy but also reduces computational complexity and enhances interpretability. In this study, we employed multiple feature selection techniques to optimize machine learning models. These methods include Variance Threshold (VTh), which removes low-variance features; SelectKBest, which ranks features based on statistical tests; and Support Vector Classifier with L1-based feature selection (SVC-L1), which leverages sparsity to identify important features. Additionally, we utilized Tree-based feature selection (Extra Tree) to assess feature importance using ensemble learning, Recursive Feature Elimination (RFE) to iteratively refine the most relevant subset, and Sequential Forward Selection (SFS) to progressively add features that contribute most to the model’s performance. By integrating these diverse approaches, we ensured a robust and comprehensive feature selection process, ultimately enhancing the efficiency and reliability of our machine learning models.

### 2.7 Protein Language Model as classifiers

Utilizing classical machine learning (ML) models laid the groundwork for our research, but we advanced our approach by incorporating pre-trained language models (PLMs). These computational frameworks leverage sophisticated natural language processing techniques to analyze the intricate details of protein structures, functions, and interactions. In particular, we employed several models from the Evolutionary Scale Modeling (esm) series, specifically esm2-t33, esm2-t30, esm2-t12, and esm2-t6. These models are pre-trained on extensive protein sequence datasets and excel in various tasks such as structure prediction and variant effect analysis. The esm series has demonstrated remarkable capabilities in capturing the evolutionary relationships and functional aspects of proteins due to their architecture and training methodology [76]. In addition to the esm models, we also integrated ProtBERT into our framework. ProtBERT is a protein-specific model derived from the BERT architecture and is pre-trained on a vast corpus of protein sequences. This model is particularly adept at understanding the contextual relationships within protein sequences, which enhances its performance in tasks related to protein classification and function prediction. The combination of these models significantly improved the accuracy and reliability of our classification system for neurotoxic and non-toxic peptides [80,81].

To fine-tune these models for our task, we optimized their hyperparameters of these models. For ProtBERT, sequences were tokenized with maximum length sequences and a batch size of 32. The model was fine-tuned across multiple epochs (6, 12, and 32) using the AdamW optimizer with a learning rate of 1X 10^-5^. Similarly, for esm, sequences were tokenized, and the *esmForSequenceClassification* model was trained for the same range of epochs using Adam with an identical learning rate. Cross-entropy loss was utilized, and both models employed truncation, padding, and gradient updates to enhance classification performance for neurotoxic and non-toxic peptides.

Overall, the integration of PLMs like esm2 and ProtBERT into our research not only enhanced our classification model’s performance but also underscored the importance of leveraging state-of-the-art computational techniques in understanding complex biological phenomena. These advancements pave the way for more accurate predictions and deeper insights into protein functionality and interactions, ultimately contributing to fields such as drug discovery and synthetic biology.

### 2.7 Cross-validation and performance metrics

As outlined in the Data Collection and Pre-processing section, for model training and evaluation, datasets were split into 80% training and 20% independent test sets. The test set remained unused during training or tuning. Peptide and protein training sets were merged into a combined training dataset, and their test sets were merged into a combined independent dataset. To prevent data leakage and ensure robust model performance estimates, a stratified five-fold cross-validation procedure was implemented exclusively within the training set. Stratified cross-validation maintains the original class proportions in each fold, ensuring that the characteristics of the original dataset are preserved during the evaluation process. This approach provides a more reliable estimate of model performance compared to simple k-fold cross-validation, especially when dealing with imbalanced datasets. The effectiveness of various machine learning models was evaluated using both threshold-dependent and threshold-independent metrics. The area under the receiver operating characteristic curve (AUC) serves as a threshold-independent measure, while sensitivity, specificity, accuracy, and the Matthews correlation coefficient (MCC) are classified as threshold-dependent metrics. These evaluation criteria have been thoroughly documented in earlier research studies [9,27].

## 3. Results

### 3.1 Compositional analysis of different toxicities

Firstly, we compare the average amino acid composition of neurotoxic and non-toxic sequences. The amino acid profiles are analyzed across all three datasets, and a comparative assessment of their average amino acid compositions was conducted (Supplementary Figure S1). Notably, the occurrence of cysteine (C), a polar and uncharged amino acid, was significantly greater in the neurotoxin sequences compared to those of non-toxins. This observation aligns with trends identified in previous research [27]. Statistical analysis identified significant differences in amino acid composition between neurotoxic and non-toxic peptides. Neurotoxic peptides exhibited a significantly higher abundance of cysteine, glycine, and lysine, with a corrected p-value of 0.001. Conversely, non-toxic peptides demonstrated a significantly higher composition of alanine (A), isoleucine (I), leucine (L), and glutamine (Q) across all three datasets.

In the past number of methods have been developed for predicting different types of toxicities. We compare composition profile of different type of toxicities to understand preference of different amino acids. In Figure 4, we show composition profile of six distinct datasets: neurotoxic peptides, neurotoxic proteins, combined neurotoxic datasets (peptides + proteins), hemotoxic peptides, toxic peptides (general toxicity), and the general genome. This aims to identify amino acid compositional biases that distinguish functionally specialized toxic biomolecules from the broader genomic background. Neurotoxic peptides and proteins are analyzed separately and collectively to evaluate whether their amino acid profiles converge or diverge, while hemotoxic and general toxic peptides are included to assess if neurotoxicity correlates with unique compositional signatures. The inclusion of the general genome provides a baseline for distinguishing natural amino acid distributions from those enriched in toxic entities. Key observations likely include elevated frequencies of hydrophobic or charged residues (e.g., cysteine, tryptophan) in neurotoxic and hemotoxic peptides, which are often critical for membrane interactions or receptor targeting. Conversely, the general genome may show higher proportions of smaller, residues (e.g., alanine, glycine). This comparative visualization highlights how evolutionary and functional pressures shape amino acid utilization in toxic molecules, offering insights into potential structural or mechanistic hallmarks of toxicity.

**Figure 4:**
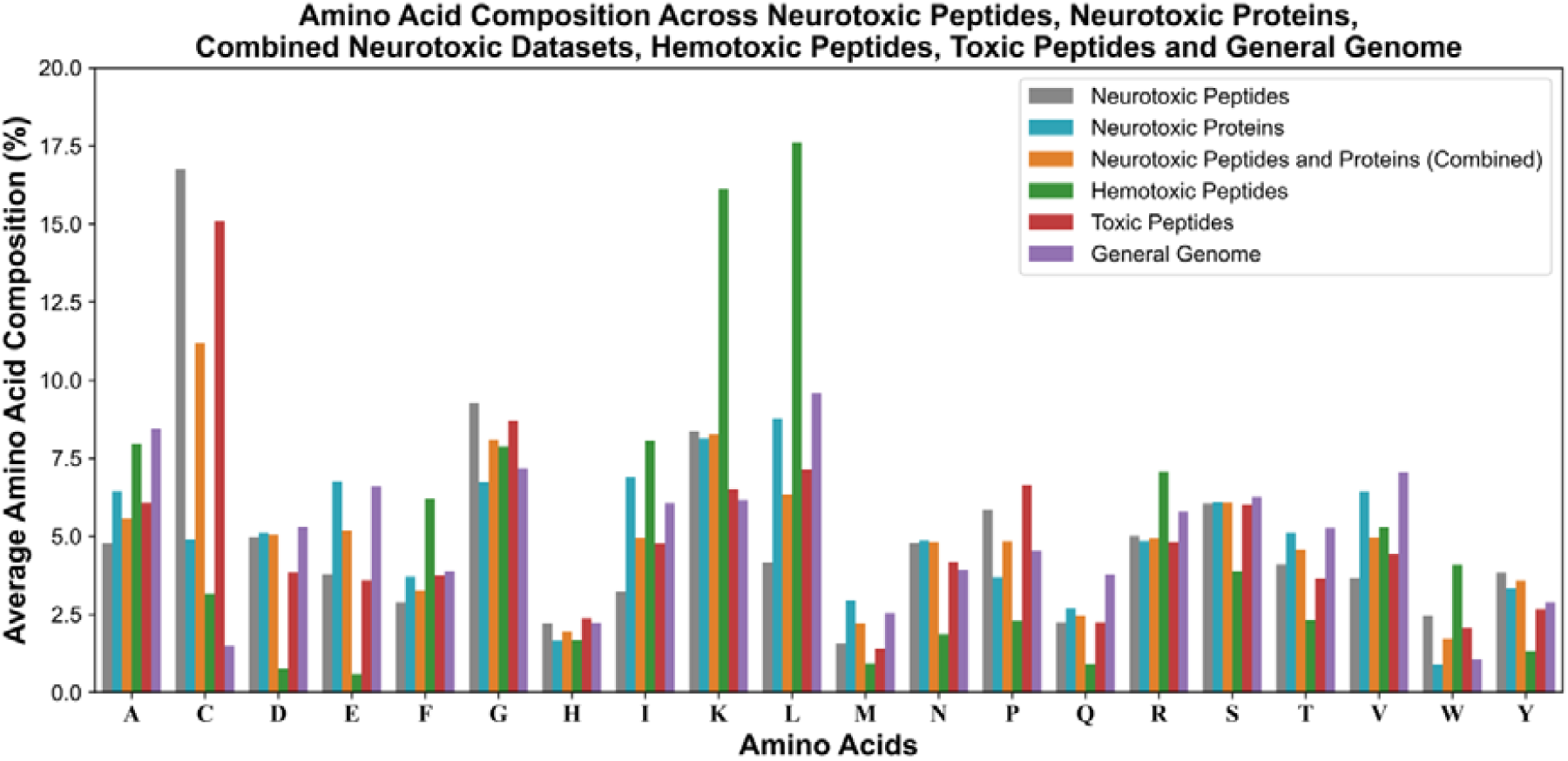
Composition profile of the different types of toxicity-associated proteins and peptides; it includes neurotoxic peptides/proteins, hemotoxic, general toxic peptides, and the human genome.

### 2.3 Correlation between toxicities

The amino acid composition correlation matrix reveals distinct relationships among various toxic components and the general genome (Table 1). It was observed that neurotoxic peptides and neurotoxins have a poor correlation that further supports the development of separate models for neurotoxic peptides and neurotoxins. Neurotoxic peptides show a strong positive correlation with the overall toxin profile (r□≈□0.94), indicating their significant role in neurotoxicity. Neurotoxic peptides have a negligible correlation with the general genome (r□≈□–0.03), suggesting a high degree of specialization, diverging from the general genome. Neurotoxic proteins exhibit moderate correlations with the general genome (r□≈□0.86), reflecting a balance between maintaining essential physiological functions and contributing to neurotoxic effects. Hemotoxins display a moderate correlation with neurotoxic proteins (r□≈□0.68) and the general genome (r□≈□0.61), but a low correlation with neurotoxic peptides (r□≈□0.18), suggesting that hemotoxins and neurotoxins may have distinct evolutionary pathways and functional roles. These findings align with research indicating that snake venoms are complex mixtures of proteins and peptides, each evolving to target specific physiological systems, such as the nervous or circulatory systems, in their prey [82].

**Table 1:** Confusion matrix shows a correlation between different types of toxicities and the general genome. Amino acid residues significantly higher (corrected p-value <0.05) in each dataset are highlighted using their single-letter codes, with color coding applied to distinguish between datasets.

|  | Neurotoxic peptides | Neurotoxic proteins | Neurotoxic combined | Hemotoxins | Toxins | General |
| --- | --- | --- | --- | --- | --- | --- |
| Neurotoxic peptides | 1 | 0.325<br>C, G, H, P, W, Y | 0.916<br>C, G, H, P, W | 0.178<br>C, D, E, G, H, M, N, P, Q, S, T, Y | 0.942<br>C, D, G, K, N, T, W, Y | -0.027<br>C, G, K, N, P, W, Y |
| Neurotoxic proteins | 0.325<br>A, E, F, I, L, M, Q, T, V | 1 | 0.678<br>A, E, F, I, L, M, Q, T, V | 0.677<br>C, D, E, M, N, P, Q, S, T, V, Y | 0.454<br>A, D, E, I, K, L, M, N, Q, T, V, Y | 0.862<br>C, I, K, M, N, Y |
| Neurotoxic combined | 0.916<br>A, E, F, I, L, M, T, V | 0.678<br>C, G, H, P, W, Y | 1 | 0.426<br>C, D, E, M, N, P, Q, S, T, Y | 0.925<br>D, E, K, M, N, Q, T, V, Y | 0.345<br>C, G, K, N, P, W, Y |
| Hemotoxins | 0.178 | 0.677 | 0.426 | 1 | 0.337 | 0.610 |
|  | A, F, I, K, L, R, V, W | A, F, G, I, K, L, R, W | A, F, I, K, L, R, W |  | A, F, I, K, L, R, V, W | C, F, G, I, K, L, R, W |
| Toxins | 0.942 | 0.454 | 0.925 | 0.337 | 1 | 0.167 |
|  | A, F, I, L, P, V | C, G, H, P, W | A, C, F, G, H, L, P, W | C, D, E, G, H, M, N, P, Q, S, T, Y |  | C, G, H, K, N, P, W |
| General | -0.027 | 0.862 | 0.345 | 0.610 | 0.167 | 1 |
|  | A, D, E, F, I, L, M, Q, R, T, V | A, D, F, G, H, L, P, Q, R, V, W | A, D, E, F, H, I, L, M, Q, R, S, T, V | D, E, H, M, N, P, Q, S, T, V, Y | A, D, E, F, I, L, M, Q, R, S, T, V, Y |  |

### 3.3 Positional analysis

We generated Two Sample Logos (TSLs) for three datasets to analyze amino acid preferences in neurotoxic and non-neurotoxic sequences, as shown in Figure 5. The TSLs highlight the positional significance of amino acid residues at a p-value threshold of 0.05. In neurotoxic peptides, cysteine (C) is predominantly favored across all positions except the first, indicating its critical role at both N- and C-termini (Figure 5A). Additionally, lysine (K) and tryptophan (PPV) are more prevalent in the C-terminal region, suggesting a preference for these residues in this segment. Non-toxic peptides predominantly feature methionine (M) at the first position and leucine (L) at the C-terminus. In contrast, neurotoxic proteins show a preference for cysteine (C) residues at positions 8 to 11 of the C-terminal region (Figure 5B). Residues like lysine (K), leucine (L), and isoleucine (I) are more favored at the N-terminal. The combined dataset exhibits a pattern similar to the peptide dataset (Figure 5C). These findings indicate distinct amino acid preferences between protein and peptide datasets, highlighting the specialized roles of specific residues in neurotoxic functions.

**Figure 5:**
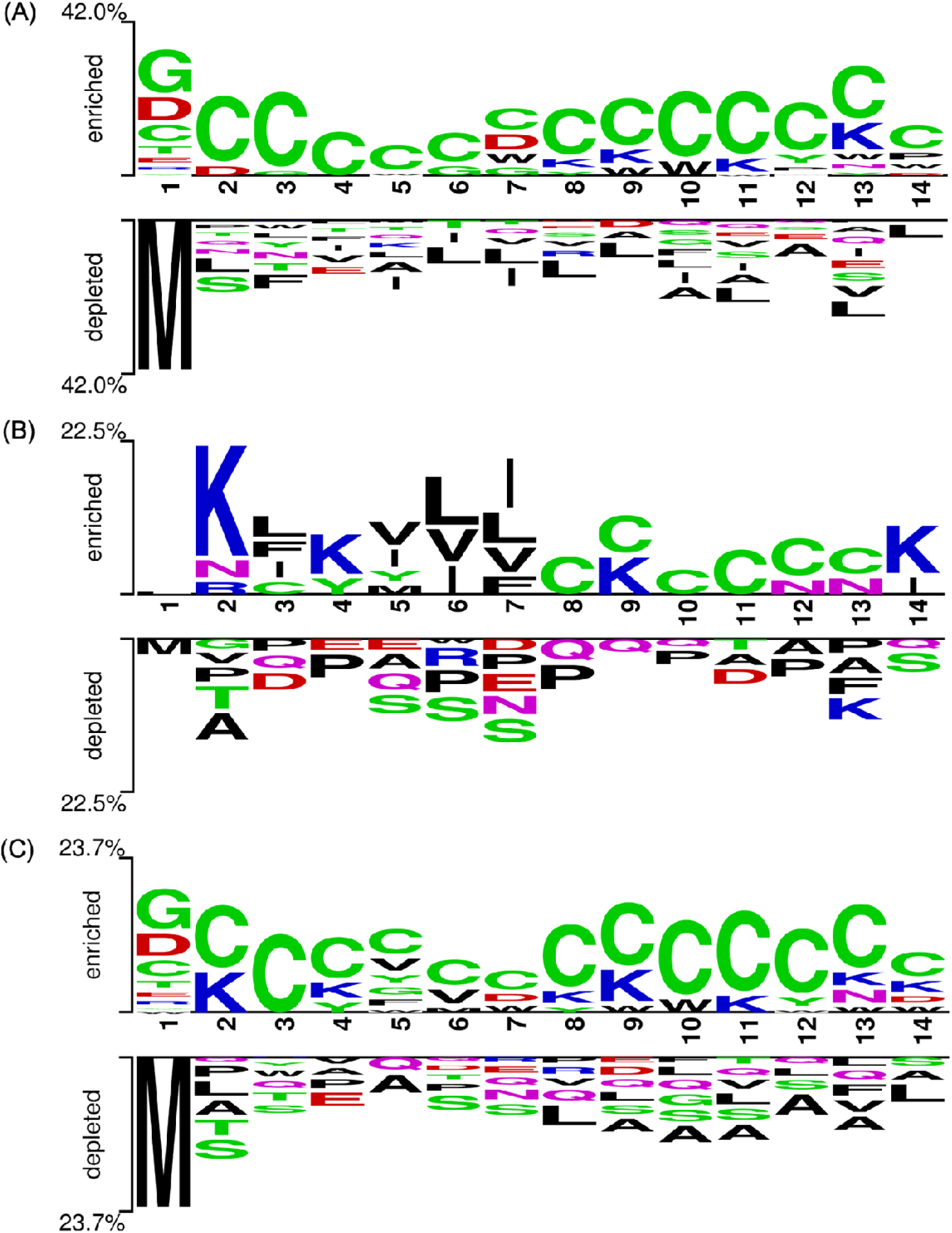
Residue preferences at different positions in neurotoxic and non-toxic sequences are illustrated using TSL for the peptide dataset (A), protein dataset (B), and combined dataset (C). The first seven positions represent the N-terminal region, while the last seven positions represent the C-terminal region of the peptides.

### 3.4 Amino Acid Composition-based Prediction

In this analysis, we evaluated the predictive power of amino acid composition for distinguishing between neurotoxic and non-toxic sequences by examining three datasets: peptide, protein, and a combined set. In Supplementary Table S4, results reveal significant differences in the amino acid profiles; for instance, cysteine (C) consistently emerged as a key differentiator. Cysteine shows markedly different mean values between neurotoxic and non-toxic peptides compared to proteins, suggesting that peptides have a unique compositional profile that may confer specific functional attributes related to neurotoxicity. In the peptide dataset, the dramatic contrast in the composition of cysteine with neurotoxic sequences exhibiting much higher values than non-toxic ones results in high accuracy (0.867) and AUC (0.901), whereas the protein dataset shows a less pronounced difference, with lower accuracy and AUC values. Similar trends are observed for other amino acids like leucine (L), where the separation between neurotoxic and non-toxic groups differs significantly between peptides and proteins. These compositional disparities underline the biological and structural differences inherent to peptides versus full-length proteins. Consequently, this suggests the development of separate predictive models for peptides and proteins, as a unified model may fail to capture the nuanced variations in amino acid distributions that are critical for accurately discriminating neurotoxins from non-neurotoxins. Overall, this compositional analysis highlights the utility of amino acid profiles as discriminative features for the prediction of neurotoxicity, offering valuable insights that may aid in understanding the molecular determinants of neurotoxic activity. A detailed results are present in Supplementary Table S4.

### 3.4 Physicochemical Property-based Prediction

Physicochemical property-based prediction analysis reveals that the predictive utility of physicochemical properties differs markedly across the peptide, protein, and combined datasets. In the peptide dataset, descriptors such as the composition of sulfur-containing residues (PCP_SC) and advanced indices like PCP_Z3 (Polarity / Charge ratio) and PCP_Z5 achieved high accuracy (0.830) and AUC (0.879), indicating these properties can strongly discriminate between neurotoxic and non-toxic peptides. In contrast, for the protein dataset, most properties showed substantially lower performance with accuracies generally around 50–60% and AUCs not exceeding 0.70–0.75, suggesting that the inherent differences in physicochemical profiles between neurotoxic and non-toxic proteins are less pronounced. The combined dataset produced intermediate metrics, reflecting a dilution of discriminative power when peptides and proteins are analyzed together. Overall, these findings critically support the development of separate, predictive models, as the features that robustly distinguish neurotoxicity in peptides do not translate equivalently to proteins. Comprehensive results are available in Supplementary Table S5.

### 3.5 Analysis of Cysteine Count

The cysteine distribution is a key determinant of peptide and protein stability, particularly in neurotoxic peptides, which predominantly feature even cysteine counts. This facilitates disulfide bond formation, enhancing structural rigidity, protease resistance, and bioactivity, as seen in conotoxins and scorpion toxins [83,84]. In peptide, dataset analysis shows that among 877 neurotoxic peptides, 738 have an even cysteine count, while only 78 have an odd count and 61 lack cysteine (Table 2). In contrast, non-toxic peptides exhibit a more balanced distribution, with 195 having even counts and 515 lacking cysteine. This suggests that even-numbered cysteines can be a defining feature of neurotoxins, stabilizing their three-dimensional conformation and enhancing toxicity. In neurotoxic proteins, the difference in even and odd cysteine counts is less pronounced; similarly, non-toxic proteins show no strong preference. The higher odd-cysteine occurrence in proteins indicates additional stabilizing factors, such as hydrogen bonding or salt bridges [85].

**Table 2:** Preference for even and odd cysteine counts in neurotoxin datasets.

| Datasets |  | Sequences without cysteine | Sequences with even cysteine count | Sequences with odd cysteine count |
| --- | --- | --- | --- | --- |
| Peptides | Neurotoxin (877) | 61 | 738 | 78 |
|  | Non-toxin (877) | 515 | 195 | 167 |
| Protein | Neurotoxin (775) | 66 | 385 | 324 |
|  | Non-toxin (775) | 186 | 247 | 342 |

### 3.5 Machine Learning-based Models

Four distinct feature sets were employed to develop machine learning models for classifying neurotoxic peptides: binary profiles, PSSM profiles, *Pfeature* compositional features, 56D *modlAMP* descriptors, and protein embeddings generated from large language models (LLM). These feature sets served as input for subsequent machine learning model training and evaluation using the Scikit-learn package [86].

#### 3.5.1 Binary Profiles

Using NT7 and CT7 binary profile features, a range of machine learning models were developed for all three datasets. Table 3 presents the performance metrics best models utilizing binary composition-based features. Among the evaluated various machine learning models, tree-based algorithms, specifically ET and RF demonstrated superior performance on all three datasets. The ET models achieved AUC values of 0.949 and 0.846 on peptide and combined independent datasets, respectively. In the protein-independent dataset, the RF model exhibited AUC values of 0.976. Similarly, using the same strategy, we also generated NT15 and CT15 binary profile features. However, the performance remained comparable. Supplementary Table S6 provides performance metrics of all the models on different datasets.

**Table 3:** Performance of best machine learning models neurotoxic datasets using binary profile features.

| Dataset | Classifier | Coss-validation Dataset |  |  |  |  |  | Independent Dataset |  |  |  |  |  |
| --- | --- | --- | --- | --- | --- | --- | --- | --- | --- | --- | --- | --- | --- |
|  |  | SENS | SPEC | PPV | ACC | MCC | AUC | SENS | SPEC | PPV | ACC | MCC | AUC |
| Peptides | RF | 0.909 | 0.864 | 0.870 | 0.887 | 0.774 | 0.945 | 0.931 | 0.841 | 0.853 | 0.886 | 0.775 | 0.946 |
|  | ET | 0.905 | 0.860 | 0.866 | 0.882 | 0.766 | 0.942 | 0.914 | 0.847 | 0.856 | 0.880 | 0.763 | 0.949 |
| Protein | RF | 0.677 | 0.706 | 0.698 | 0.692 | 0.384 | 0.752 | 0.665 | 0.716 | 0.701 | 0.690 | 0.381 | 0.776 |
|  | Lasso | 0.661 | 0.703 | 0.690 | 0.682 | 0.365 | 0.743 | 0.658 | 0.755 | 0.729 | 0.706 | 0.415 | 0.756 |
| Combined | ET | 0.804 | 0.727 | 0.747 | 0.766 | 0.533 | 0.836 | 0.797 | 0.731 | 0.747 | 0.764 | 0.529 | 0.846 |
|  | RF | 0.830 | 0.692 | 0.729 | 0.761 | 0.527 | 0.830 | 0.833 | 0.683 | 0.724 | 0.758 | 0.522 | 0.841 |

SENS = sensitivity; SPEC = specificity; PPV = precision; ACC = accuracy; AUC = area under receiver operating characteristic; MCC = Matthews correlation coefficient

#### 3.5.2 PSSM Profiles

This study investigates the role of evolutionary information in predicting neurotoxins. Due to the short length of peptides, their PSSM profiles are unreliable, so we extracted PSSM features only for protein dataset. Among various PSSM-derived features, PSSM composition performed best, achieving an AUC of 0.889 with the ET model on independent protein dataset. When combined with AAC composition, the performance varied *aac_pssm* + AAC composition improved the AUC to 0.861, whereas combining *pssm_composition* or *mepd_pssm* with AAC composition led to a decline in performance (Table 4). These findings highlight the selective impact of evolutionary features on neurotoxin prediction. Supplementary Table S7 provides detailed performance metrics of all the models on different datasets.

**Table 4:** The performance of best machine learning models on protein dataset using binary profile features and combination with AAC composition features.

| Features | Classifier | Coss-validation Dataset |  |  |  |  |  | Independent Dataset |  |  |  |  |  |
| --- | --- | --- | --- | --- | --- | --- | --- | --- | --- | --- | --- | --- | --- |
|  |  | SENS | SPEC | PPV | ACC | MCC | AUC | SENS | SPEC | PPV | ACC | MCC | AUC |
| aac_pssm (20) | SVC (rbf) | 0.785 | 0.757 | 0.765 | 0.771 | 0.543 | 0.835 | 0.781 | 0.742 | 0.752 | 0.761 | 0.523 | 0.841 |
| aac_pssm (20) + AAC composition (20) | SVC (rbf) | 0.814 | 0.802 | 0.804 | 0.808 | 0.616 | 0.871 | 0.800 | 0.787 | 0.790 | 0.794 | 0.587 | 0.861 |
| pssm_composition (400) | ET | **0.837** | **0.832** | **0.833** | **0.834** | **0.669** | **0.885** | **0.813** | **0.806** | **0.808** | **0.810** | **0.619** | **0.889** |
| pssm_composition (400) + AAC composition (20) | ET | 0.672 | 0.716 | 0.702 | 0.694 | 0.388 | 0.760 | 0.697 | 0.710 | 0.706 | 0.703 | 0.406 | 0.747 |
| mepd_pssm (420) | RF | 0.768 | 0.772 | 0.771 | 0.770 | 0.540 | 0.841 | 0.806 | 0.735 | 0.753 | 0.771 | 0.543 | 0.851 |
| mepd_pssm (420) + AAC composition (20) | EF | 0.735 | 0.727 | 0.729 | 0.731 | 0.462 | 0.797 | 0.748 | 0.639 | 0.674 | 0.694 | 0.389 | 0.766 |

(SENS: Sensitivity; SPEC: Specificity; PPV: Positive Predictive Value; ACC: Accuracy; MCC: Matthews Correlation Coefficient; AUC: Area Under the Receiver Operating Characteristic Curve; The bold values indicate the best-performing models for each dataset)

#### 3.5.3 Compositional Features

Using *Pfeature*, 9190 features were generated per sequence and used to train and evaluate machine-learning models. Among the evaluated models, tree-based models consistently exhibited superior predictive performance. Table 5 summarizes the performance of the best-performing models on all three datasets across different feature combinations. The RF model achieved a peak AUC of 0.969 and 0.873 on peptide and protein-independent datasets, respectively, using ALLCOMP features. While ALLCOMP features provided the highest accuracy, their utilization resulted in a significant computational burden. The prediction time for the full ALLCOMP set was approximately 515.23 seconds. In contrast, employing AAC and DPC features reduced the computational time to 48.51 seconds with only a minor decline in AUC. All computational experiments were conducted on a high-performance computing system featuring an Intel(R) Xeon(R) Gold 6338 CPU (2.00GHz, 16 cores, 64-bit), 100.58 GB of RAM, and 768 MB L3 cache, operating within a VMware virtualization environment. In the case of a combined independent dataset, the ET model performs best with an AUC of 0.919 on AAC features.

**Table 5:**
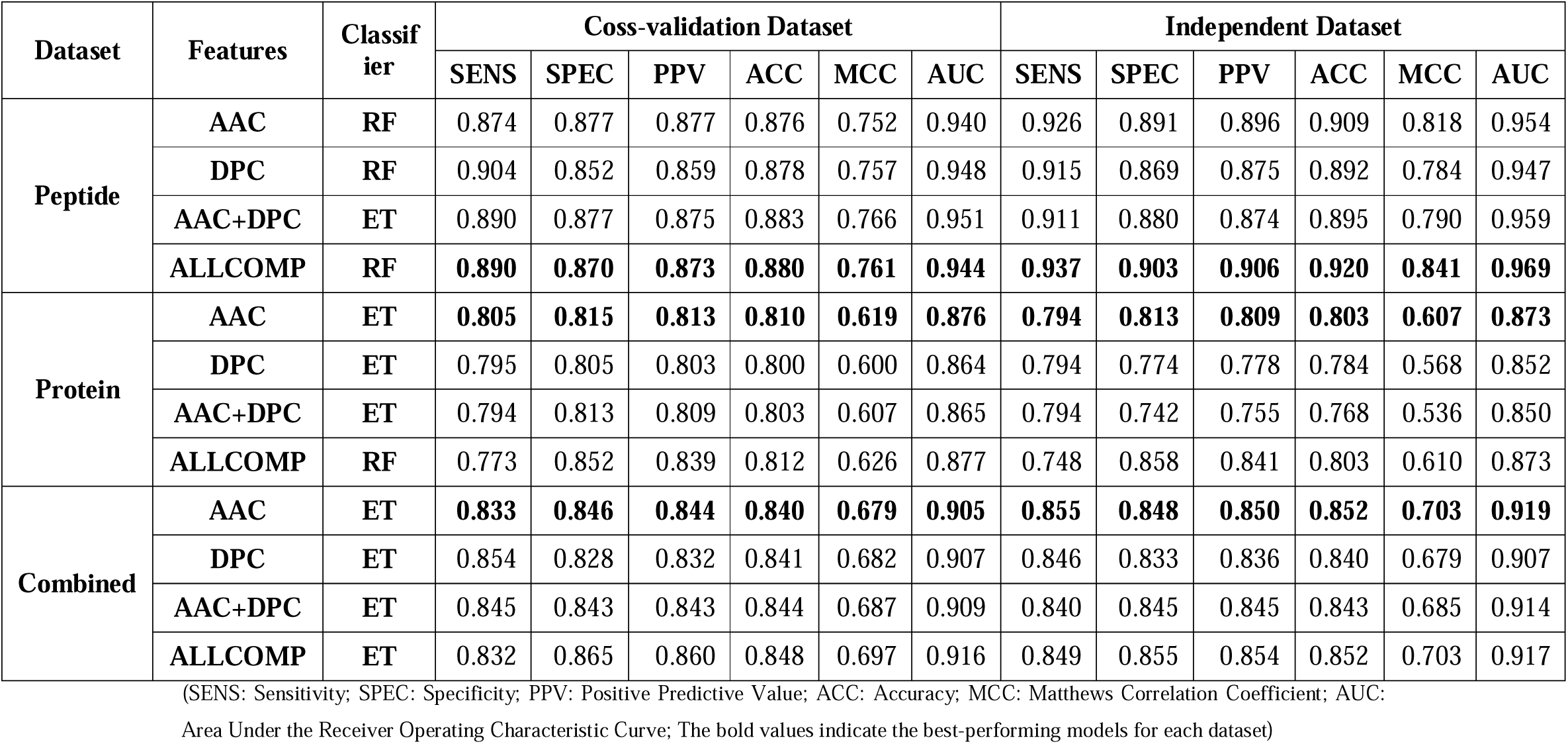
The performance of best machine learning models developed using different compositional features on all datasets.

Similarly, the *modlAMP* library is used to compute sequence-based features, such as physicochemical and structural properties, and facilitates the development of machine learning models for neurotoxic peptide classification and prediction. On the peptide dataset, the ET model exhibited superior performance among other models, achieving an AUC of 0.934 on independent data, as detailed in Supplementary Table S8. On a protein and combined dataset, the RF model demonstrated best performance, attaining an AUC of 0.845 and 0.907 on protein and combined independent dataset, respectively.

#### 3.5.4 Embeddings from Fine-tuned PLMs

Embedding-based machine learning models have become increasingly important in various domains due to their ability to effectively represent complex data types in a lower-dimensional space [87–89]. In this study, we utilized embeddings from various fine-tuned esm checkpoints and the ProtBERT model. These embeddings, derived from fine-tuned protein language models (PLMs) such as esm and ProtBERT, capture both evolutionary and structural information. On the peptide dataset, ProtBERT embeddings using the LR model achieved the best performance with an AUC of 0.963 on independent data. For the protein and combined datasets, esm2-t30 embeddings with the ET model performed best, attaining an AUC of 0.934 and 0.949 on protein and combined independent data, respectively. Detailed results are presented in Table 6.

**Table 6:** The performance of machine learning models developed using PLM embeddings on all three datasets.

| Dataset | Embedding Source | Classifier | Coss-validation Data |  |  |  |  |  | Independent Data |  |  |  |  |  |
| --- | --- | --- | --- | --- | --- | --- | --- | --- | --- | --- | --- | --- | --- | --- |
|  |  |  | SENS | SPEC | PPV | ACC | MCC | AUC | SENS | SPEC | PPV | ACC | MCC | AUC |
| Peptide | esm2-t6 | ET | 0.882 | 0.883 | 0.883 | 0.882 | 0.765 | 0.944 | 0.897 | 0.858 | 0.863 | 0.877 | 0.756 | 0.947 |
|  | esm2-t12 | ET | 0.869 | 0.902 | 0.898 | 0.885 | 0.771 | 0.949 | 0.880 | 0.881 | 0.880 | 0.880 | 0.761 | 0.941 |
|  | esm2-t30 | ET | 0.875 | 0.914 | 0.911 | 0.895 | 0.790 | 0.955 | 0.880 | 0.886 | 0.885 | 0.883 | 0.766 | 0.954 |
|  | esm2-t33 | ET | 0.870 | 0.920 | 0.916 | 0.895 | 0.791 | 0.951 | 0.891 | 0.903 | 0.902 | 0.897 | 0.795 | 0.950 |
|  | esm2-t36 | RF | 0.859 | 0.913 | 0.908 | 0.886 | 0.773 | 0.952 | 0.863 | 0.915 | 0.910 | 0.889 | 0.779 | 0.953 |
|  | ProtBERT | LR | **0.932** | **0.919** | **0.920** | **0.925** | **0.850** | **0.964** | **0.920** | **0.891** | **0.895** | **0.906** | **0.812** | **0.963** |
| Protein | esm2-t6 | ET | 0.803 | 0.860 | 0.851 | 0.831 | 0.664 | 0.902 | 0.800 | 0.865 | 0.855 | 0.832 | 0.666 | 0.896 |
|  | esm2-t12 | ET | 0.790 | 0.874 | 0.863 | 0.832 | 0.667 | 0.908 | 0.787 | 0.910 | 0.897 | 0.848 | 0.702 | 0.918 |
|  | esm2-t30 | ET | **0.832** | **0.894** | **0.887** | **0.863** | **0.727** | **0.923** | **0.819** | **0.903** | **0.894** | **0.861** | **0.725** | **0.934** |
|  | esm2-t33 | ET | 0.817 | 0.905 | 0.906 | 0.851 | 0.725 | 0.925 | 0.806 | 0.931 | 0.924 | 0.844 | 0.737 | 0.930 |
|  | esm2-t36 | ET | 0.806 | 0.923 | 0.912 | 0.865 | 0.734 | 0.922 | 0.800 | 0.918 | 0.919 | 0.854 | 0.717 | 0.924 |
|  | ProtBERT | ET | 0.727 | 0.815 | 0.797 | 0.771 | 0.544 | 0.853 | 0.729 | 0.774 | 0.764 | 0.752 | 0.504 | 0.861 |
| Combined | esm2-t6 | ET | 0.832 | 0.878 | 0.872 | 0.855 | 0.711 | 0.920 | 0.831 | 0.888 | 0.881 | 0.859 | 0.720 | 0.931 |
|  | esm2-t12 | ET | 0.831 | 0.888 | 0.881 | 0.860 | 0.720 | 0.927 | 0.840 | 0.906 | 0.900 | 0.873 | 0.748 | 0.934 |
|  | esm2-t30 | ET | **0.846** | **0.899** | **0.893** | **0.872** | **0.746** | **0.936** | **0.855** | **0.930** | **0.925** | **0.893** | **0.787** | **0.949** |
|  | esm2-t33 | SVC (linear) | 0.839 | 0.864 | 0.864 | 0.866 | 0.733 | 0.931 | 0.850 | 0.912 | 0.909 | 0.891 | 0.773 | 0.949 |
|  | esm2-t36 | ElasticNet | 0.871 | 0.886 | 0.884 | 0.879 | 0.757 | 0.945 | 0.870 | 0.876 | 0.875 | 0.873 | 0.746 | 0.943 |
|  | ProtBERT | ET | 0.795 | 0.803 | 0.801 | 0.799 | 0.597 | 0.886 | 0.831 | 0.803 | 0.809 | 0.817 | 0.634 | 0.903 |

(SENS: Sensitivity; SPEC: Specificity; PPV: Positive Predictive Value; ACC: Accuracy; MCC: Matthews Correlation Coefficient; AUC: Area Under the Receiver Operating Characteristic Curve; The bold values indicate the best-performing models for each dataset)

#### 3.5.5 Feature Selection

The evaluation of various feature selection techniques across peptide, protein, and combined datasets (Table 7) highlights the potential of dimensionality reduction in maintaining or enhancing predictive performance. In the peptide dataset, we applied various feature selection methods to the ALLCOMP features, reducing them to subsets such as SVCL1 (467 features) and tree-based selection (100 features), which achieved comparable AUCs of 0.955 and 0.952, respectively. Notably, the ET model based on simple AAC+DPC features outperformed the feature selection-based approaches. In contrast, feature selection proved highly effective for the protein and combined datasets. In the protein dataset, RFE using only 48 features achieved an independent AUC of 0.937, demonstrating that a compact feature subset is sufficient for robust neurotoxin prediction. Similarly, in the combined dataset, the SVCL1 method, which selected just 84 features, attained the highest independent AUC of 0.954. These findings underscore the importance of feature selection in reducing model complexity while preserving or even enhancing predictive accuracy, particularly in protein and combined datasets. Techniques such as RFE and SVCL1 effectively reduced the number of features while maintaining or improving key performance indicators, including AUC, sensitivity, and MCC, making them valuable strategies for optimizing neurotoxicity prediction models.

**Table 7:** The performance of extra tree model using different feature selection methods across datasets.

| Dataset | Featutre Selection method | No of features | Coss-validation Data |  |  |  |  |  | Independent Data |  |  |  |  |  |
| --- | --- | --- | --- | --- | --- | --- | --- | --- | --- | --- | --- | --- | --- | --- |
|  |  |  | SENS | SPEC | PREC | ACC | MCC | AUC | SENS | SPEC | PREC | ACC | MCC | AUC |
| Peptides | VTh | 184 | 0.892 | 0.883 | 0.884 | 0.887 | 0.775 | 0.950 | 0.926 | 0.886 | 0.891 | 0.906 | 0.813 | 0.956 |
|  | k-best | 55 | 0.890 | 0.872 | 0.874 | 0.881 | 0.762 | 0.947 | 0.920 | 0.880 | 0.885 | 0.900 | 0.801 | 0.954 |
|  | RFE | 148 | 0.876 | 0.866 | 0.867 | 0.871 | 0.742 | 0.945 | 0.920 | 0.891 | 0.895 | 0.906 | 0.812 | 0.956 |
|  | SFS | 128 | 0.874 | 0.870 | 0.871 | 0.872 | 0.745 | 0.941 | 0.920 | 0.886 | 0.890 | 0.903 | 0.807 | 0.944 |
|  | SVCL1 | 467 | 0.906 | 0.885 | 0.887 | 0.895 | 0.791 | 0.959 | 0.909 | 0.897 | 0.899 | 0.903 | 0.806 | 0.955 |
|  | Tree based | 100 | 0.894 | 0.872 | 0.874 | 0.883 | 0.766 | 0.946 | 0.915 | 0.880 | 0.885 | 0.897 | 0.795 | 0.952 |
|  | No Feature Selection | 420<br>(AAC+DPC) | 0.890 | 0.877 | 0.875 | 0.883 | 0.766 | 0.951 | 0.911 | 0.880 | 0.874 | 0.895 | 0.790 | 0.959 |
| Proteins | VTh | 67 | 0.821 | 0.895 | 0.887 | 0.858 | 0.718 | 0.914 | 0.813 | 0.903 | 0.894 | 0.858 | 0.719 | 0.935 |
|  | k-best | 50 | 0.824 | 0.861 | 0.856 | 0.843 | 0.686 | 0.917 | 0.826 | 0.877 | 0.871 | 0.852 | 0.704 | 0.929 |
|  | RFE | 48 | 0.850 | 0.889 | 0.884 | 0.869 | 0.739 | 0.929 | 0.826 | 0.903 | 0.895 | 0.865 | 0.731 | 0.937 |
|  | SFS | 50 | 0.794 | 0.877 | 0.866 | 0.835 | 0.673 | 0.905 | 0.794 | 0.884 | 0.872 | 0.839 | 0.680 | 0.924 |
|  | SVCL1 | 66 | 0.840 | 0.881 | 0.876 | 0.860 | 0.722 | 0.927 | 0.852 | 0.877 | 0.874 | 0.865 | 0.729 | 0.934 |
|  | Tree based | 65 | 0.831 | 0.871 | 0.866 | 0.851 | 0.702 | 0.919 | 0.826 | 0.884 | 0.877 | 0.855 | 0.711 | 0.930 |
|  | No Feature Selection (640 esm2-t30 embeddings) | 640 | 0.832 | 0.894 | 0.887 | 0.863 | 0.727 | 0.923 | 0.819 | 0.903 | 0.894 | 0.861 | 0.725 | 0.934 |
| Combined | VTh | 66 | 0.833 | 0.893 | 0.886 | 0.863 | 0.728 | 0.926 | 0.849 | 0.915 | 0.909 | 0.882 | 0.766 | 0.941 |
|  | k-best | 50 | 0.830 | 0.886 | 0.879 | 0.858 | 0.717 | 0.926 | 0.825 | 0.894 | 0.886 | 0.859 | 0.720 | 0.943 |
|  | RFE | 70 | 0.853 | 0.896 | 0.892 | 0.875 | 0.750 | 0.937 | 0.855 | 0.927 | 0.922 | 0.891 | 0.784 | 0.949 |
|  | SFS | 50 | 0.839 | 0.890 | 0.884 | 0.864 | 0.729 | 0.924 | 0.843 | 0.903 | 0.897 | 0.873 | 0.747 | 0.930 |
|  | SVCL1 | 84 | 0.852 | 0.898 | 0.893 | 0.875 | 0.751 | 0.940 | 0.870 | 0.909 | 0.906 | 0.890 | 0.780 | 0.954 |
|  | Tree based | 50 | 0.835 | 0.891 | 0.885 | 0.863 | 0.727 | 0.927 | 0.846 | 0.903 | 0.897 | 0.874 | 0.750 | 0.940 |
|  | No Feature Selection (640 esm2-t30 embeddings) | 640 | 0.846 | 0.899 | 0.893 | 0.872 | 0.746 | 0.936 | 0.855 | 0.93 | 0.925 | 0.893 | 0.787 | 0.949 |

(SENS: Sensitivity; SPEC: Specificity; PPV: Positive Predictive Value; ACC: Accuracy; MCC: Matthews Correlation Coefficient; AUC: Area Under the Receiver Operating Characteristic Curve; The bold values indicate the best-performing models for each dataset)

### 3.6 Protein Language Models

In this work, we employed various PLMs to predict neurotoxic proteins and peptides. Since these models are not explicitly trained or fine-tuned for classifying neurotoxic peptides, they primarily rely on the general representations learned from large, diverse datasets. To tailor these models for the specific task, we optimized their hyperparameters using a training dataset composed of neurotoxic and non-toxic proteins and peptides, enabling them to perform effectively in this specialized classification task. The performance of fine-tuned protein language models on the different datasets is shown in Table 8. In the peptide dataset among all models, the esm2-t30 achieved the best performance on the independent data with an AUC of 0.984, while the esm2-t6 excelled on the protein and combined independent data with an AUC of 0.912 and 0.943, respectively.

**Table 8:** The performance evaluation of PLMs on all the datasets.

| Dataset | PLM Classifier | Coss-validation Data |  |  |  |  |  | Independent Data |  |  |  |  |  |
| --- | --- | --- | --- | --- | --- | --- | --- | --- | --- | --- | --- | --- | --- |
|  |  | SENS | SPEC | PPV | ACC | MCC | AUC | SENS | SPEC | PPV | ACC | MCC | AUC |
| Peptides | esm2-t6 | 0.892 | 0.881 | 0.882 | 0.923 | 0.770 | 0.952 | 0.960 | 0.847 | 0.862 | 0.903 | 0.817 | 0.950 |
|  | esm2-t12 | 0.879 | 0.901 | 0.882 | 0.883 | 0.776 | 0.951 | 0.942 | 0.887 | 0.862 | 0.913 | 0.820 | 0.961 |
|  | esm2-t30 | **0.937** | **0.941** | **0.944** | **0.943** | **0.877** | **0.981** | **0.926** | **0.970** | **0.970** | **0.951** | **0.898** | **0.984** |
|  | esm2-t33 | 0.648 | 0.972 | 0.835 | 0.937 | 0.712 | 0.952 | 0.952 | 0.880 | 0.883 | 0.911 | 0.830 | 0.961 |
|  | ProtBERT | 0.976 | 0.862 | 0.876 | 0.919 | 0.843 | 0.970 | 0.949 | 0.800 | 0.827 | 0.875 | 0.758 | 0.961 |
| Proteins | esm2-t6 | **0.89** | **0.903** | **0.917** | **0.891** | **0.800** | **0.930** | **0.804** | **0.890** | **0.903** | **0.841** | **0.693** | **0.912** |
|  | esm2-t12 | 0.907 | 0.898 | 0.911 | 0.894 | 0.843 | 0.940 | 0.839 | 0.787 | 0.797 | 0.810 | 0.630 | 0.900 |
|  | esm2-t30 | 0.888 | 0.912 | 0.910 | 0.900 | 0.801 | 0.953 | 0.804 | 0.873 | 0.864 | 0.838 | 0.678 | 0.912 |
|  | esm2-t33 | 0.834 | 0.870 | 0.865 | 0.852 | 0.704 | 0.915 | 0.833 | 0.859 | 0.855 | 0.846 | 0.692 | 0.905 |
|  | ProtBERT | 0.827 | 0.800 | 0.805 | 0.813 | 0.627 | 0.877 | 0.813 | 0.791 | 0.796 | 0.802 | 0.604 | 0.875 |
| Combined | esm2-t6 | **0.921** | **0.886** | **0.889** | **0.900** | **0.803** | **0.952** | **0.867** | **0.875** | **0.875** | **0.873** | **0.742** | **0.943** |
|  | esm2-t12 | 0.872 | 0.869 | 0.854 | 0.897 | 0.793 | 0.941 | 0.778 | 0.894 | 0.893 | 0.84 | 0.68 | 0.922 |
|  | esm2-t30 | 0.853 | 0.856 | 0.856 | 0.855 | 0.710 | 0.917 | 0.865 | 0.865 | 0.865 | 0.865 | 0.729 | 0.931 |
|  | esm2-t33 | 0.848 | 0.860 | 0.858 | 0.854 | 0.708 | 0.907 | 0.865 | 0.871 | 0.870 | 0.868 | 0.735 | 0.920 |
|  | ProtBERT | 0.777 | 0.815 | 0.807 | 0.796 | 0.592 | 0.868 | 0.755 | 0.787 | 0.780 | 0.771 | 0.542 | 0.846 |

(SENS: Sensitivity; SPEC: Specificity; PPV: Positive Predictive Value; ACC: Accuracy; MCC: Matthews Correlation Coefficient; AUC: Area Under the Receiver Operating Characteristic Curve; The bold values indicate the best-performing models for each dataset)

### 3.7 Final Models

In this study, we developed a range of ML models and protein language models to predict neurotoxic and non-toxic peptides and proteins using three distinct datasets. Among these, the ESM2-t30 model demonstrated the highest performance on the peptide dataset, achieving an AUC of 0.984 and an MCC of 0.898. For both the protein and combined datasets, the ET model, utilizing ESM2-t30 embeddings as features, exhibited superior predictive capability. In the protein dataset, the ET model achieved the best performance with an AUC of 0.937 and an MCC of 0.731, having been trained on 48 embeddings selected through RFE. Similarly, in the combined dataset, the ET model attained optimal performance with an AUC of 0.954 and an MCC of 0.780, leveraging 84 embeddings selected via the SVCL1 method. These findings highlight the effectiveness of feature selection techniques in optimizing model performance across different datasets.

### 3.8 Cross Dataset Prediction of Final Models

In this section, we evaluated the performance of the models across all three datasets to determine their generalizability. Specifically, we assessed whether a model trained on peptides could effectively predict neurotoxic proteins, whether a model trained on proteins could accurately predict neurotoxic peptides, and whether a combined model exhibited superior predictive performance compared to models trained exclusively on peptide or protein datasets. This analysis provides insights into whether separate models are necessary for peptides and proteins or if a unified model can effectively generalize across both categories.

In Table 9 cross-dataset analysis reveals significant variations in model performance depending on the dataset used for training and validation. The ESM2-t30 model, developed on the neurotoxic peptide dataset, exhibits exceptional performance on the independent peptide dataset (AUCLJLJ0.984), but its perform very poor on protein dataset (AUC 0.752). This discrepancy suggests that the peptide-specific model is not well-suited for identifying neurotoxic proteins, likely due to intrinsic differences in sequence composition and structural features between peptides and proteins. ET model developed on the neurotoxic protein dataset shows high performance on the independent protein dataset (AUCLJLJ0.937) but perform poorly on peptide dataset (AUC 0.883). Notably, the ET model developed on the combined dataset delivers consistently high performance across all independent datasets, achieving AUCs of 0.954, 0.939, and 0.967 for the combined, protein, and peptide datasets, respectively. However, the method developed on combined dataset perform poorer than model trained on peptides for predicting peptides. Overall, it is clear from above analysis that there is need to develop separate model for proteins and peptides.

**Table 9.** Cross-Dataset Performance of Final Models.

| Final Model | Dataset | SENS | SPEC | PPV | ACC | MCC | AUC |
| --- | --- | --- | --- | --- | --- | --- | --- |
| **ESM2-t30**<br>**(peptide**<br>**Dataset)** | **Independent**<br>**peptide dataset** | **0.926** | **0.970** | **0.970** | **0.951** | **0.898** | **0.984** |
|  | **Independent**<br>**protein dataset** | 0.386 | 0.981 | 0.951 | 0.674 | 0.441 | 0.752 |
|  | **Independent**<br>**combined dataset** | 0.665 | 0.976 | 0.963 | 0.821 | 0.674 | 0.879 |
| **Extra Tree**<br>**(Protein**<br>**Dataset)** | **Independent**<br>**protein dataset** | **0.826** | **0.903** | **0.895** | **0.865** | **0.731** | **0.937** |
|  | **Independent**<br>**peptide dataset** | 0.653 | 0.931 | 0.906 | 0.792 | 0.608 | 0.883 |
|  | **Independent**<br>**combined dataset** | 0.734 | 0.918 | 0.9 | 0.826 | 0.663 | 0.904 |
| **Extra Tree**<br>**(combined**<br>**Dataset)** | **Independent**<br>**combined dataset** | **0.870** | **0.909** | **0.906** | **0.890** | **0.780** | **0.954** |
|  | **Independent**<br>**protein dataset** | 0.838 | 0.91 | 0.903 | 0.874 | 0.75 | 0.939 |
|  | **Independent**<br>**peptide dataset** | 0.892 | 0.914 | 0.912 | 0.903 | 0.807 | 0.967 |

(SENS: Sensitivity; SPEC: Specificity; PPV: Positive Predictive Value; ACC: Accuracy; MCC: Matthews Correlation Coefficient; AUC: Area Under the Receiver Operating Characteristic Curve; The bold values indicate the best-performing models in particular dataset)

### 3.9 Benchmarking with Existing Methods

Thorough comparing against existing methods is crucial for evaluating the advancements of newly developed approaches. NTxPred2, an updated version of NTxPred, incorporates several key innovations, including the integration of fine-tuned protein language models (PLMs), novel composition and physicochemical features. Table 10 presents a comparative analysis of the proposed NTxPred2 framework against its predecessor, NTxPred, using independent datasets. To ensure a comprehensive evaluation, NTxPred2 was benchmarked against various neurotoxicity prediction models previously developed by Saha et al. Among these, the AAC-based SVM model exhibited the highest performance on the independent peptide dataset, achieving an AUC of 0.922. For protein and combined independent datasets, the SVM (DPC) model demonstrated superior performance, with AUC values of 0.715 and 0.783, respectively. Additionally, the CNN model developed by Lee et al. (2021) achieved an AUC of 0.689 on the independent protein dataset. However, a direct comparative evaluation with tools developed by Yang et al. (2008) [33], Guang et al. (2010) [30], Song et al. (2012) [34], Tang et al. (2017) [35], Huo et al. (2017) [36], Koua et al. (2017) [32], Mei et al. (2018) [37], and Wan et al. (2023) [40] was not feasible due to the unavailability of their underlying algorithms. Pippin, a random forest-based method [38], could not be compared due to the web service being nonfunctional. The results clearly demonstrate that NTxPred2 outperforms existing methods. Its consistently superior performance highlights its potential as a valuable tool for therapeutic peptide development, particularly in the classification of neurotoxic proteins and peptides.

**Table 10:** Benchmarking of Performance of existing methods on independent datasets of NTxPred2.

| Tool | Model | Data | SENS | SPEC | PPV | ACC | MCC | AUC |
| --- | --- | --- | --- | --- | --- | --- | --- | --- |
| NTxPred | SVM (DPC + Length) | Independent peptide dataset | 0.790 | 0.794 | 0.794 | 0.792 | 0.584 | 0.827 |
|  |  | Independent protein dataset | 0.412 | 0.761 | 0.634 | 0.587 | 0.185 | 0.589 |
|  |  | Independent combined dataset | 0.589 | 0.603 | 0.598 | 0.596 | 0.192 | 0.656 |
|  | SVM (DPC) | Independent peptide dataset | 0.898 | 0.811 | 0.827 | 0.855 | 0.712 | 0.916 |
|  |  | Independent protein dataset | 0.639 | 0.619 | 0.627 | 0.629 | 0.258 | 0.715 |
|  |  | Independent combined dataset | 0.652 | 0.772 | 0.742 | 0.712 | 0.428 | 0.783 |
|  | SVM (AAC + Length) | Independent peptide dataset | 0.813 | 0.709 | 0.737 | 0.761 | 0.524 | 0.832 |
|  |  | Independent protein dataset | 0.529 | 0.658 | 0.607 | 0.593 | 0.188 | 0.648 |
|  |  | Independent combined dataset | 0.607 | 0.755 | 0.712 | 0.680 | 0.365 | 0.692 |
|  | SVM (AAC) | Independent peptide dataset | 0.932 | 0.777 | 0.808 | 0.855 | 0.718 | 0.922 |
|  |  | **Independent protein dataset** | 0.574 | 0.651 | 0.622 | 0.612 | 0.226 | 0.669 |
|  |  | **Independent combined dataset** | 0.661 | 0.766 | 0.640 | 0.714 | 0.430 | 0.770 |
| Lee et al. (2021) | CNN | **Independent peptide dataset** | 0.00 | 1.00 | 0.00 | 0.498 | 0.00 | 0.500 |
|  |  | **Independent protein dataset** | 0.277 | 0.993 | 0.977 | 0.635 | 0.388 | 0.635 |
|  |  | **Independent combined dataset** | 0.130 | 0.996 | 0.977 | 0.563 | 0.245 | 0.563 |
| NTxPred2 | **ESM2-t30 (Peptide Dataset)** | **Independent peptide dataset** | **0.926** | **0.970** | **0.970** | **0.951** | **0.898** | **0.984** |
|  | **Extra Tree (Protein Dataset)** | **Independent protein dataset** | **0.826** | **0.903** | **0.895** | **0.865** | **0.731** | **0.937** |
|  | **Extra Tree (Combined Dataset)** | **Independent combined dataset** | **0.870** | **0.909** | **0.906** | **0.890** | **0.780** | **0.954** |

(SENS: Sensitivity; SPEC: Specificity; PPV: Positive Predictive Value; ACC: Accuracy; MCC: Matthews Correlation Coefficient; AUC: Area Under the Receiver Operating Characteristic Curve)

### 3.10 Design and Deployment of a Web Server

To enhance the accessibility and utility of our research findings, we developed NTxPred2, a comprehensive platform consisting of both a user-friendly web server, standalone software, and *pip* package. This platform is freely accessible to the scientific community. The NTxPred2 web server offers a suite of functionalities, including peptide and protein neurotoxicity prediction, protein scanning for potential neurotoxic regions, and designing of non-toxic peptides/proteins. For users requiring high-throughput analysis, a standalone software and *pip* package is available for download, enabling efficient and large-scale neurotoxic peptide prediction.

## 4. Discussion

The genomes of numerous organisms have been sequenced over the past three decades due advancements in sequencing technology. This has resulted in a rapid expansion of protein databases, which store information on protein sequences, structures, functions, and therapeutic applications. A major challenge in the post-genomic era is the annotation of these rapidly growing databases of proteins. Highlighting the significance of protein annotation, the 2024 Nobel Prize was awarded to researchers for their groundbreaking contributions to the structural annotation of proteins. Therapeutic annotation of a protein is another crucial aspect of annotation, where researchers are exploiting therapeutic applications of proteins. Over the past few decades, the US FDA has approved a significant number of proteins for therapeutic applications, recognizing their potential in treating various diseases [90–92]. In addition to sequencing, genetically modified organisms, genetically modified crops, and recombinant technology have played a key role in the large-scale production of therapeutic proteins like monoclonal antibodies and hormones [93,94]. The technological advancement has also played a crucial role in shifting vaccine development from traditional whole-organism vaccines to protein-based vaccines [95,96].

In summary, the use of proteins in therapeutics is rapidly increasing, presenting significant challenges in assessing their safety for humans, animals, and the environment. To address these concerns, the World Health Organization (WHO) has established guidelines for evaluating the safety of genetically modified (GM) foods and protein-based therapies. Toxicity assessment is a crucial component of the safety evaluation of genetically modified (GM) foods and therapeutic proteins. Ensuring that these proteins do not cause adverse effects is essential for their regulatory approval and clinical use. Previously, a number of computational tools have been developed to predict and assess cytotoxicity, hemotoxicity, immunotoxicity, and neurotoxicity. In this study, we made a systematic attempt to predict the neurotoxicity of proteins, a critical factor that often leads to “black box warnings” on medications, as emphasized by Walker et al. [97,98].

Our primary analysis revealed significant differences in the amino acid composition of neurotoxic sequences compared to non-toxic sequences, particularly a higher frequency of cysteine. Enrichment of cysteine in neurotoxins enables disulfide-bonded scaffolds, conferring protease resistance and structural stability, a hallmark of venom peptides evolved under predator-prey arms races [99]. We also compare amino acid composition of neurotoxic peptides and neurotoxins and observed significance difference. Neurotoxic peptides exhibit a very high association with overall toxin content, reflecting rapid divergence driven by strong positive selection for optimized physiochemical properties such as charge and hydrophobicity that enhance ion channel interactions [100,101]. In contrast, neurotoxin typically larger and preserving conserved residue like inhibitory cystine knots display a strong correlation to the general genome, suggesting that their evolutionary trajectory is tempered by the need to maintain structural integrity while permitting functional diversification [102,103]. This composition difference justify our hypothesis to develop separate method for predicting neurotoxic peptides and for predicting neurotoxins. We also compare amino acid composition of proteins/peptides having different type of toxicities. Our primary analysis justify development model for predicting neurotoxic peptides/proteins.

Composition and physicochemical property-based prediction analysis highlight significant differences in discriminatory power between neurotoxins and non-toxins. In peptides, cystine and leucine composition alone achieved high predictive performance, with AUC values of 0.90 and 0.75 and accuracies of 0.87 and 0.70, respectively. However, in proteins, cystine-based composition performed less effectively, with an AUC of 0.69 and an accuracy of 0.65. Physicochemical descriptors such as polarity (PCP_PO), non-polar residues (PCP_NP), aliphatic side chains (PCP_AL), and sulfur-containing residues (PCP_SC) demonstrated strong predictive power in peptides but were less effective in proteins. Notably, advanced indices (PCP_Z3, PCP_Z5) remained reliable across both datasets, indicating their robustness in toxicity prediction. Finally, the cysteine count analysis revealed that non-toxic peptides tend to favor odd cysteine counts, whereas neurotoxic peptides are more likely to exhibit even cysteine counts. This finding suggests that for the development of therapeutic peptides, particularly those intended to be non-toxic, careful consideration should be given to cysteine distribution, as odd cysteine counts may contribute to reduced toxicity.

This study explored a diverse range of feature-generation and feature-selection strategies to optimize predictive performance. Among the traditional machine learning models, tree based models (ET and RF) exhibited superior performance. In peptide datasets, the esm2-t30 model performed best (AUC 0.98) on peptide-independent datasets. For both the protein and combined datasets, the ET model, utilizing ESM2-t30 embeddings as features, exhibited superior predictive capability. In the protein dataset, the ET model achieved the best performance with an AUC of 0.94 and an MCC of 0.73, having been trained on 48 embeddings selected through RFE. Similarly, in the combined dataset, the ET model attained optimal performance with an AUC of 0.95 and an MCC of 0.78, using 84 embeddings selected via the SVCL1 method. Cross-dataset predictions demonstrated superior performance for models trained on the peptide dataset compared to those trained on combined or protein datasets. While the protein model (ET) achieved an AUC of 0.937, comparable to the combined model (AUC 0.939), these results underscore inherent compositional and functional differences between peptides and proteins, supporting the development of specialized models. Although the combined model integrates diverse sequence characteristics, the observed performance differences suggest that domain-specific models remain preferable for maximizing predictive accuracy within their respective classes.

### Future Directions

The increasing need for accurate toxicity assessment in therapeutic proteins and GM foods calls for a unified platform capable of predicting all types of toxicity, including allergenicity/ immunogenicity, and off-target effects. Such a system should adhere to WHO guidelines, ensuring comprehensive safety evaluations covering toxicity, allergenicity, gene stability, and unintended effects. Additionally, minimizing reliance on animal testing is crucial, as traditional in vivo models are time-consuming, ethically challenging, and limited in scalability. In silico tools offer a powerful alternative by using computational algorithms to predict toxicity with high accuracy, aligning with the 3Rs principle (Replacement, Reduction, and Refinement) in research. Developing and refining such computational models will accelerate drug discovery, improve safety assessments, and support regulatory decision-making, ultimately leading to safer and more effective therapeutic interventions.

### Conclusion

This study introduces NTxPred2, a comprehensive platform for classifying neurotoxic and non-toxic sequences. NTxPred2 integrates a high-accuracy classification model and is freely accessible via a user-friendly web server. All three best-performing models are deployed on our server. To cater to diverse research needs, a pip package and a standalone software package are also provided. While NTxPred2 offers significant advancements in neurotoxicity prediction, it is important to acknowledge certain limitations. Currently, the platform relies solely on peptide sequence information for classification and does not incorporate information regarding the peptide’s source of origin and structure. Furthermore, NTxPred2 is primarily designed for natural peptides, excluding non-canonical amino acids, modified peptides, and peptides shorter than seven residues. This focused approach, while enhancing model efficiency, may limit the applicability of NTxPred2 to specific peptide classes.

## Authors’ contributions

ASR collected and processed the datasets. ASR and SJ implemented the algorithms and developed the prediction models. SJ created the front-end user interface, ASR created the back end of the web server, and SC analyzed the results. ASR, SC, and GPSR penned the manuscript. GPSR conceived and coordinated the project. All authors have read and approved the final manuscript.

## Acknowledgments

The authors express their gratitude to the University Grants Commission (UGC), Council of Scientific and Industrial Research (CSIR), Department of Science and Technology (DST-INSPIRE), and Department of Biotechnology (DBT) for their generous fellowships and financial support. They also thank the Department of Computational Biology, IIITD, New Delhi, for the excellent infrastructure and facilities. The authors would like to acknowledge the Department of Biotechnology (DBT) for the infrastructure grant awarded to the institute. Furthermore, they would like to acknowledge BioRender.com for creating the figures utilized in this work. The authors acknowledge the use of ChatGPT in improving the language of the manuscript.

## Data Availability

The datasets generated for this study can be accessed on the ‘NTxPred2’ web server at https://webs.iiitd.edu.in/raghava/ntxpred2/download.php and publicly available on GitHub https://github.com/raghavagps/ntxpred2.

## Code availability

The source code for this study is publicly available on GitHub and can be found at https://github.com/raghavagps/ntxpred2 and the ‘NTxPred2’ web server at https://webs.iiitd.edu.in/raghava/ntxpred2/download.php.

