## Supplementary Data for "NTxPred2: A large language model for predicting neurotoxic peptides and neurotoxins"

**Mailing Address of Authors**

**^*^Corresponding Author**

Prof. Gajendra P. S. Raghava

Head and Professor

Department of Computational Biology

Indraprastha Institute of Information Technology, Delhi

Okhla Industrial Estate, Phase III, (Near Govind Puri Metro Station)

New Delhi, India – 110020

Office: A-302 (R&D Block)

Website: <http://webs.iiitd.edu.in/raghava/>

| Table | Content | Page No. |
| --- | --- | --- |
| *Supplementary Table S1* | Overview of Computational tools for predicting different types of toxic activities of peptides and proteins. | 3 |
| *Supplementary Table S2* | List of all the composition-based features computed from Pfeature standalone along with vector length. | 6 |
| *Supplementary Table S3* | List of Physicochemical descriptors extracted using modlAMP. | 7 |
| *Supplementary Figure S1* | Average amino acid composition of neurotoxin and non-toxin sequences in different datasets. | 8 |
| *Supplementary Table S4* | Amino Acid Composition-based Prediction. | 9 |
| *Supplementary Table S5* | Physicochemical Property-based Prediction -based Prediction. | 12 |
| *Supplementary Table S6* | Performance evaluation of machine learning models on the different datasets using binary profile features. | 16 |
| *Supplementary Table S7* | Performance evaluation of machine learning models on the protein datasets using PSSM profiles. | 19 |
| *Supplementary Table S8* | Performance of best machine learning models on neurotoxic datasets using modlAMP features and combination with compositional features. | 23 |

Supplementary Table S1: Overview of Computational tools for predicting different types of toxic activities of peptides and proteins.

**Supplementary Table S1: Overview of computational tools for predicting different types of toxic activities of peptides and proteins.**

| **Tool (year)** | **Description (link)** | **Reference** |
| --- | --- | --- |
| **General Toxicity Prediction Tools** | | |
| BTXpred (2007) | Classification of bacterial toxins  (https://webs.iiitd.edu.in/raghava/btxpred/) | [5] |
| ToxinPred (2014) | Toxicity prediction of peptides and proteins  (https://webs.iiitd.edu.in/raghava/toxinpred/) | [49] |
| SpiderP (2013) | Propeptide cleavage sites prediction in spider toxins  (http://PPV.arachnoserver.org/spiderP.html) | [50] |
| ToxClassifier (2016) | Venom toxins prediction  (http://bioserv7.bioinfo.pbf.hr/ToxClassifier/) | [12] |
| TOXIFY (2019) | Prediction of animal venom proteins using deep learning (https://PPV.github.com/tijeco/toxify) | [13] |
| ATSE (2021) | Structural and evolutionary information based toxicity prediction (http://server.malab.cn/ATSE) | [15] |
| ToxIBLT (2022) | A deep learning based method for toxicity prediction of peptide (http://server.wei-group.net/ToxIBTL) | [16] |
| ToxinPred2.0 (2022) | A method for predicting toxins  (https://webs.iiitd.edu.in/raghava/toxinpred2/) | [11] |
| ToxinPred3.0 (2024) | Updated version of ToxinPred for predicting toxicity of peptides (https://webs.iiitd.edu.in/raghava/toxinpred3/) | [9] |
| VISH-pred (2024) | Toxin prediction using LLM (http://ec2-35-170-123-194.compute-1.amazonaws.com:7860/) | [19] |
| MultiToxPred 1.0 | Prediction and classification of protein toxins (https://www.biochemintelli.com/MultiToxPred-v1) | [20] |
| **Hemolytic Toxicity Prediction Tools** | | |
| HemoPI (2016) | Website and mobile application designed for computing the hemolytic activity of peptides. (https://webs.iiitd.edu.in/raghava/hemopi/) | [41] |
| HemoPred (2017) | RF-based Web server dedicated to predicting the hemolytic potency of peptides. (http://codes.bio/hemopred/) | [42] |
| HemoPImod (2020) | A web-based prediction model for chemically modified hemolytic peptides. (https://webs.iiitd.edu.in/raghava/hemopimod/) | [43] |
| HLPpred-Fuse (2020) | Enhanced and resilient prediction of hemolytic peptide activity through the fusion of multiple feature representations. (http://thegleelab.org/HLPpred-Fuse) | [42] |
| HAPPENN (2020) | An innovative tool for predicting hemolytic potency in therapeutic peptides utilizing neural networks. (https://research.timmons.eu/happenn) | [44] |
| AMPDeep (2022) | PLMs based hemolytic activity prediction of antimicrobial peptides (https://zenodo.org/records/6992526) | [45] |
| Ansari et al. (2023) | Serverless prediction of peptide properties using recurrent neural networks, focusing on classification tasks such as hemolysis, nonfouling, and solubility. (https://peptide.bio) | [46] |
| PeptideBERT (2023) | A transformer-based language model for peptide property prediction, specializing in classification tasks such as hemolysis, nonfouling, and solubility. (https://github.com/ChakradharG/PeptideBERT) | [47] |
| HemoPI2 (2024) | Prediction of Hemolytic Peptides and their Hemolytic Concentration (HC_50_) (https://webs.iiitd.edu.in/raghava/hemopi2/) | [8] |
| **Immunotoxicity/Allergenicity Prediction Tools** | | |
| Stadler et al. | Motif based Allergenicity prediction by protein sequence. | [51] |
| AlgPred (2006) | prediction of allergenic proteins and mapping of IgE epitopes. (https://webs.iiitd.edu.in/raghava/algpred/) | [52] |
| AllerHunter (2009) | SVM-based allergenicity and cross-reactivity prediction of Proteins. (http://tiger.dbs.nus.edu.sg/AllerHunter) | [53] |
| ProAP (2013) | sequence-based (ProAp-SVM), motif-based (ProAp-motif) and SVM-based allergen prediction (http://gmobl.sjtu.edu.cn/proAP/main.html) | [54] |
| AllerTOP (2013) | First alignment-free server for in silico prediction of allergens. (http://PPV.pharmfac.net/allertop) | [55] |
| AllergenFP (2014) | allergenicity prediction by descriptor fingerprints of proteins. (http://ddg-pharmfac.net/Allergen FP) | [56] |
| AllerCatPro (2019) | Prediction of allergenic potential of proteins based on 3D protein structure similarity and amino acid sequence. (https://allercatpro.bii.a-star.edu.sg/) | [57] |
| AlgPred2.0 (2021) | A method for predicting allergenic proteins and mapping of IgE epitopes. (https://webs.iiitd.edu.in/raghava/algpred2/) | [21] |
| AllerCatPro 2.0 (2022) | Tool to predict protein allergenicity potential. (https://allercatpro.bii.a-star.edu.sg/help.html) | [58] |
| ALLERDET (2022) | Deep Learning combination with the Decision Tree method for the prediction of allergenicity. ([http://allerdet.frangam.com](http://allerdet.frangam.com/)) | [59] |
| SEP-AlgPro (2024) | prediction tool utilizing traditional ML and DL techniques with protein language model features. (<https://balalab-skku.org/SEP-AlgPro/>) | [60] |
| **Neurotoxicity Prediction Tools** | | |
| NTxPred (2007) | Tool for predicting neurotoxins and classifying them based on their function and origin. (https://webs.iiitd.edu.in/raghava/ntxpred2/) | [27] |
| Yang et al. (2008) | Prediction of presynaptic and postsynaptic neurotoxins by the increment of diversity. | [33] |
| Guang et al. (2010) | SVM based ML model to predict neurotoxicity. | [30] |
| Song et al (2012) | Prediction of presynaptic and postsynaptic neurotoxins by bi-layer SVM with multi-features | [34] |
| Tang et al. (2017) | Predicting Presynaptic and Postsynaptic Neurotoxins by  Developing Feature Selection Technique. | [35] |
| Huo et al. (2017) | Prediction of presynaptic and postsynaptic neurotoxins by combining various Chou's pseudo components. | [36] |
| Koua et al. (2017) | Spider Neurotoxins, Short Linear Cationic Peptides and Venom Protein Classification Improved by an Automated Competition between Exhaustive Profile HMM Classifiers | [32] |
| Mei et al. (2018) | Analysis and prediction of presynaptic and postsynaptic neurotoxins by Chou’s general pseudo amino acid composition and motif features. | [37] |
| Li et al. (2020) | Pippin: A random forest-based method for identifying presynaptic and postsynaptic neurotoxins. (http://flagship.erc.monash.edu/pippin/) | [38] |
| Lee et al. (2021) | A Deep Learning Approach with Data Augmentation to Predict Novel Spider Neurotoxic Peptides. (https://github.com/bzlee-bio/NT_estimation) | [39] |
| Wan et al. (2023) | Utilize a few features to classify presynaptic and postsynaptic neurotoxins. | [40] |

Supplementary Table S2: List of all the composition-based features computed from Pfeature standalone along with vector length.

| Name of the Feature | Feature vector length |
| --- | --- |
| Amino acid composition (AAC) | 20 |
| Dipeptide composition (DPC) | 400 |
| Tripeptide composition (TPC) | 8000 |
| Atom composition (ATC) | 5 |
| Bond composition (BTC) | 4 |
| Physicochemical Properties Composition (PCP) | 30 |
| Residue Repeat Information (RRI) | 20 |
| Property Repeat Information (PRI) | 25 |
| Distance distribution of residue (DDOR) | 20 |
| Shannon-Entropy of Protein (SEP) | 1 |
| Shannon Entropy of Physicochemical Property (SPC) | 25 |
| Shannon Entropy of Residues (SER) | 20 |
| Pseudo amino acid composition (PAAC) | 21 |
| Amphiphilic pseudo amino acid composition (APAAC) | 23 |
| Quasi-sequence order (QSO) | 42 |
| Sequence Order Coupling Number (SOCN) | 2 |
| Conjoint Triad Calculation (CTC) | 343 |
| Composition enhanced Transition Distribution (CTD) | 189 |
| Total | 9190 |

Supplementary Table S3: List of Physicochemical descriptors extracted using modlAMP.

| **Descriptor** | **Description** |
| --- | --- |
| **InstabilityIndex** | Protein stability based on amino acid composition. |
| **Length** | Peptide length. |
| **MW** | Molecular weight. |
| **NetCharge** | Overall charge. |
| **IsoelectricPoint** | Isoelectric point. |
| **Aromaticity** | Relative frequency of aromatic residues (Phe, Trp, Tyr). |
| **AliphaticIndex** | Measure of protein thermal stability based on aliphatic amino acids. |
| **BomanIndex** | Potential for protein-protein interactions. |
| **HydrophobicRatio** | Frequency of hydrophobic residues (A, C, F, I, L, M, V). |
| **S_AASI/uS_AASI** | Amino acid selectivity index scale for antimicrobial peptides. |
| **modlabs_ABHPRK** | Modlabs physicochemical feature scale (Acidic, Basic, Hydrophobic, Polar, Aromatic, Kink-inducer). |
| **H_argos/uH_argos** | Hydrophobicity index using Argos amino acid scale. |
| **B_Bulkiness/uB_Bulkiness** | Amino acid side chain bulkiness scale. |
| **charge_phys** | Amino acid charge at pH 7.0 with His charge adjusted by +0.1. |
| **charge_acid** | Amino acid charge at acidic pH with His charge adjusted by +1.0. |
| **H_Eisenberg/uH_Eisenberg** | Eisenberg hydrophobicity consensus scale. |
| **Ez** | Energy of insertion of amino acid side chains into lipid bilayers. |
| **flexibility/u_flexibility** | Amino acid side chain flexibility scale. |
| **Grantham** | Scale combining side chain composition, polarity, and molecular volume. |
| **H_GRAVY/uH_GRAVY** | GRAVY hydrophobicity amino acid scale. |
| **H_HoppWoods/uH_HoppWoods** | Hopp-Woods hydrophobicity scale. |
| **ISAECI** | Isotropic Surface Area and Electronic Charge Index of amino acid side chains. |
| **H_Janin/uH_Janin** | Janin hydrophobicity scale. |
| **H_KyteDoolittle/uH_KyteDoolittle** | Kyte & Doolittle hydrophobicity scale. |
| **F_Levitt/uF_Levitt** | Levitt alpha-helix propensity scale. |
| **MSS_shape/u_MSS_shape** | Topological shape and size index of amino acid side chains. |
| **MSW** | Scale derived from PCA of molecular surface WHIM descriptors. |
| **pepArc** | Modlabs pharmacophoric feature scale (hydrophobicity, polarity, charge, proline). |
| **pepcats** | Modlabs PEPCATS pharmacophoric scale. |
| **polarity/u_polarity** | Amino acid polarity scale. |
| **PPCALI** | Modlabs PCA-derived amino acid property scale. |
| **Refractivity/u_refractivity** | Relative amino acid refractivity values. |
| **t_scale** | PCA-derived scale using GRID program probes. |
| **TM_tend/u_TM_tend** | Amino acid transmembrane propensity scale. |
| **Z3_1, Z3_2, Z3_3** | Original three-dimensional Z-scale. |
| **Z5_1, Z5_2, Z5_3, Z5_4, Z5_5** | Extended five-dimensional Z-scale. |

Supplementary Figure S1: Average amino acid composition of neurotoxin and non-toxin sequences in different datasets.

**Supplementary Figure S1.1: Average amino acid composition of neurotoxin and non-toxin peptide sequences in peptide dataset.**

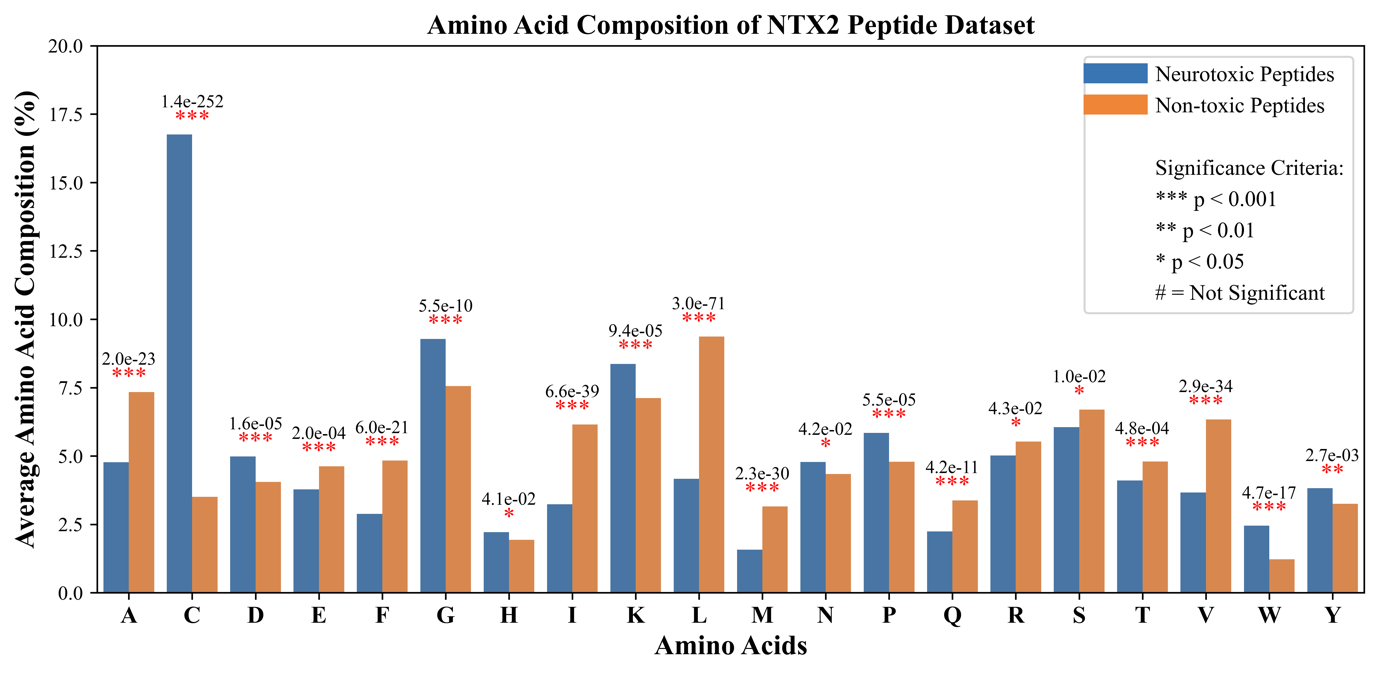

**Supplementary Figure S1.2: Average amino acid composition of neurotoxin and non-toxin protein sequences in protein dataset.**

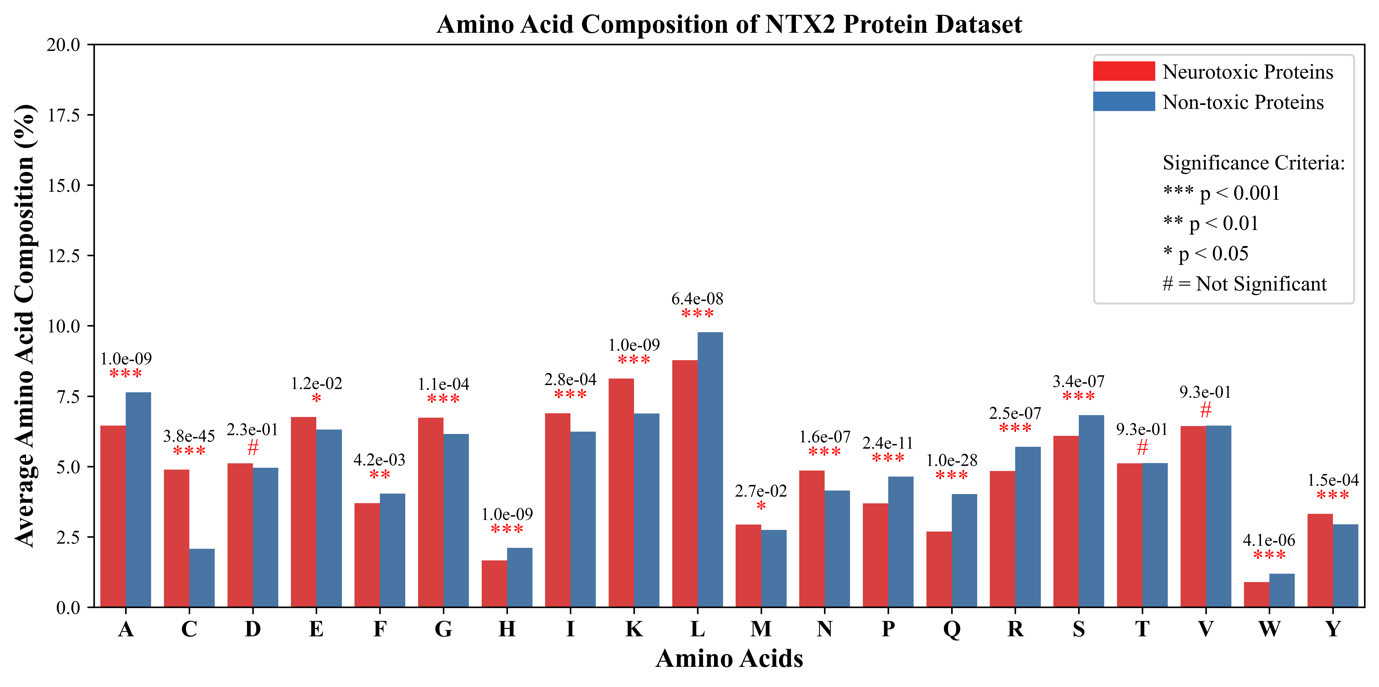

**Supplementary Figure S1.3: Average amino acid composition of neurotoxin and non-toxin peptide and protein sequences in combined dataset.**

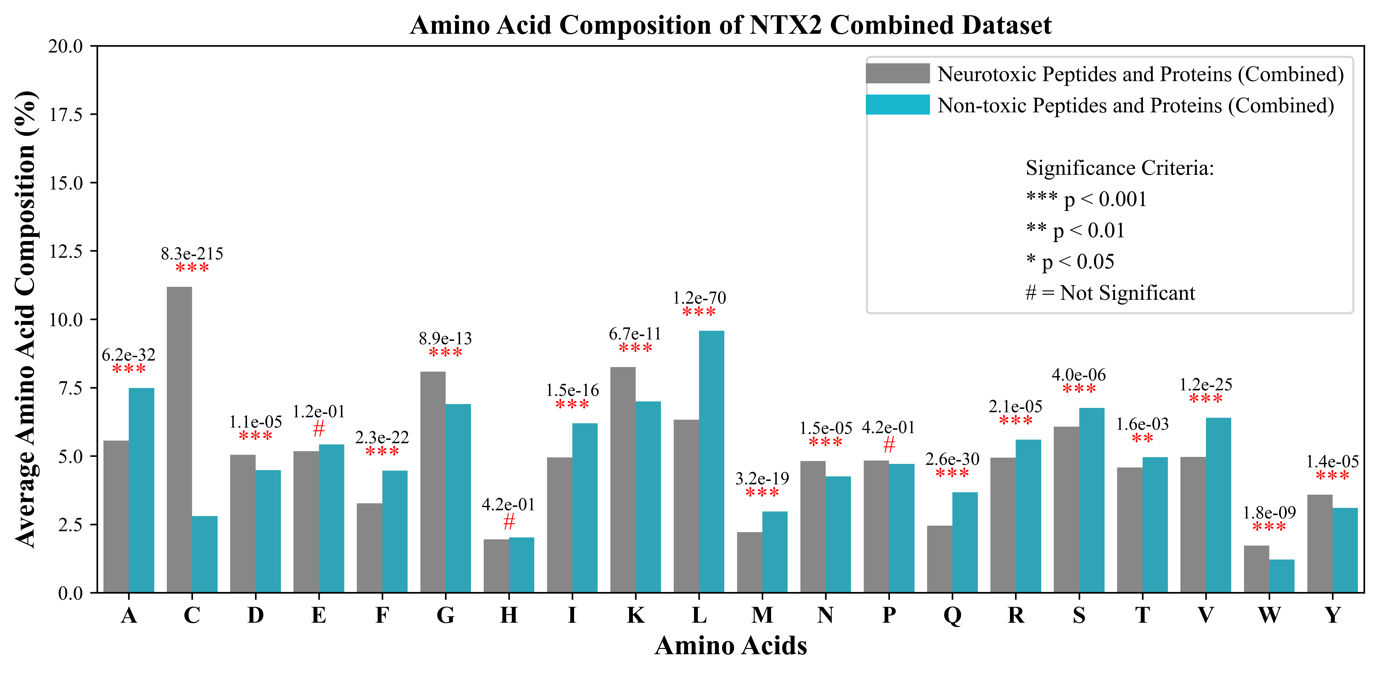

Supplementary Table S4: Amino Acid Composition-based Prediction.

**Supplementary Table S4.1: Amino acid composition-based prediction of neurotoxic and non-toxic sequences in peptide dataset.**

| Amino Acid | Best Threshold | Mean (Neurotoxic) | Mean (Non-toxic) | ACC | AUC |
| --- | --- | --- | --- | --- | --- |
| A | 6.067 | 4.773 | 7.362 | 0.604 | 0.633 |
| C | 10.103 | 16.754 | 3.452 | 0.867 | 0.901 |
| D | 4.518 | 4.979 | 4.058 | 0.544 | 0.570 |
| E | 4.207 | 3.782 | 4.631 | 0.548 | 0.564 |
| F | 3.865 | 2.884 | 4.846 | 0.609 | 0.631 |
| G | 8.418 | 9.280 | 7.555 | 0.587 | 0.616 |
| H | 2.077 | 2.214 | 1.940 | 0.543 | 0.541 |
| I | 4.694 | 3.233 | 6.156 | 0.637 | 0.676 |
| K | 7.732 | 8.370 | 7.093 | 0.547 | 0.564 |
| L | 6.785 | 4.165 | 9.405 | 0.696 | 0.746 |
| M | 2.365 | 1.572 | 3.158 | 0.616 | 0.656 |
| N | 4.560 | 4.778 | 4.343 | 0.535 | 0.540 |
| P | 5.313 | 5.846 | 4.780 | 0.560 | 0.578 |
| Q | 2.809 | 2.245 | 3.374 | 0.559 | 0.575 |
| R | 5.264 | 5.019 | 5.509 | 0.505 | 0.501 |
| S | 6.377 | 6.057 | 6.698 | 0.525 | 0.526 |
| T | 4.454 | 4.101 | 4.808 | 0.532 | 0.543 |
| V | 5.012 | 3.668 | 6.356 | 0.631 | 0.663 |
| W | 1.842 | 2.454 | 1.230 | 0.586 | 0.599 |
| Y | 3.535 | 3.823 | 3.246 | 0.533 | 0.549 |

**Supplementary Table S4.2: Amino acid composition-based prediction of neurotoxic and non-toxic sequences in the protein dataset.**

| Amino Acid | Best Threshold | Mean (Neurotoxic) | Mean (Non-toxic) | ACC | AUC |
| --- | --- | --- | --- | --- | --- |
| A | 7.046 | 6.454 | 7.637 | 0.574 | 0.584 |
| C | 3.482 | 4.889 | 2.074 | 0.645 | 0.687 |
| D | 5.037 | 5.115 | 4.960 | 0.534 | 0.536 |
| E | 6.534 | 6.757 | 6.310 | 0.543 | 0.545 |
| F | 3.871 | 3.702 | 4.041 | 0.519 | 0.529 |
| G | 6.447 | 6.735 | 6.160 | 0.543 | 0.570 |
| H | 1.887 | 1.662 | 2.112 | 0.573 | 0.593 |
| I | 6.568 | 6.896 | 6.240 | 0.557 | 0.558 |
| K | 7.505 | 8.125 | 6.885 | 0.605 | 0.610 |
| L | 9.275 | 8.782 | 9.768 | 0.575 | 0.580 |
| M | 2.844 | 2.937 | 2.751 | 0.538 | 0.554 |
| N | 4.502 | 4.855 | 4.148 | 0.577 | 0.599 |
| P | 4.167 | 3.694 | 4.641 | 0.599 | 0.601 |
| Q | 3.358 | 2.693 | 4.023 | 0.628 | 0.668 |
| R | 5.273 | 4.843 | 5.702 | 0.561 | 0.578 |
| S | 6.459 | 6.088 | 6.830 | 0.559 | 0.570 |
| T | 5.120 | 5.114 | 5.125 | 0.495 | 0.496 |
| V | 6.448 | 6.441 | 6.454 | 0.507 | 0.492 |
| W | 1.043 | 0.896 | 1.191 | 0.564 | 0.580 |
| Y | 3.133 | 3.320 | 2.947 | 0.545 | 0.562 |

**Supplementary Table S4.3: Amino acid composition-based prediction of neurotoxic and non-toxic sequences in the combined dataset.**

| Amino Acid | Best Threshold | Mean (Neurotoxic) | Mean (Non-toxic) | ACC | AUC |
| --- | --- | --- | --- | --- | --- |
| A | 6.526 | 5.561 | 7.491 | 0.588 | 0.612 |
| C | 6.997 | 11.188 | 2.806 | 0.762 | 0.808 |
| D | 4.762 | 5.043 | 4.481 | 0.537 | 0.554 |
| E | 5.298 | 5.178 | 5.419 | 0.507 | 0.518 |
| F | 3.868 | 3.268 | 4.468 | 0.567 | 0.592 |
| G | 7.493 | 8.086 | 6.900 | 0.574 | 0.599 |
| H | 1.988 | 1.955 | 2.021 | 0.515 | 0.503 |
| I | 5.573 | 4.952 | 6.195 | 0.556 | 0.582 |
| K | 7.625 | 8.255 | 6.996 | 0.569 | 0.581 |
| L | 7.953 | 6.331 | 9.575 | 0.616 | 0.674 |
| M | 2.590 | 2.212 | 2.967 | 0.545 | 0.584 |
| N | 4.533 | 4.814 | 4.251 | 0.556 | 0.563 |
| P | 4.776 | 4.837 | 4.715 | 0.485 | 0.510 |
| Q | 3.067 | 2.455 | 3.678 | 0.591 | 0.607 |
| R | 5.268 | 4.937 | 5.600 | 0.531 | 0.531 |
| S | 6.416 | 6.072 | 6.760 | 0.541 | 0.543 |
| T | 4.767 | 4.576 | 4.957 | 0.515 | 0.526 |
| V | 5.685 | 4.969 | 6.402 | 0.570 | 0.597 |
| W | 1.467 | 1.723 | 1.211 | 0.525 | 0.534 |
| Y | 3.346 | 3.587 | 3.106 | 0.540 | 0.553 |

Supplementary Table S5: Physicochemical Property-based Prediction -based Prediction.

**Supplementary Table S5.1: Physicochemical Property-based Prediction-based prediction of neurotoxic and non-toxic sequences in peptide dataset.**

| Feature | Description | Best Threshold | Mean (Neurotoxic) | Mean (Non-toxic) | Accuracy | AUC |
| --- | --- | --- | --- | --- | --- | --- |
| PCP_PC | Composition of positively charged residues | 0.151 | 0.156 | 0.146 | 0.532 | 0.554 |
| PCP_NC | Composition of negatively charged residues | 0.087 | 0.088 | 0.087 | 0.519 | 0.513 |
| PCP_NE | Composition of neutral charged residues | 0.762 | 0.756 | 0.767 | 0.531 | 0.541 |
| PCP_PO | Composition of polar residues | 0.273 | 0.330 | 0.216 | 0.737 | 0.802 |
| PCP_NP | Composition of non-polar residues | 0.443 | 0.379 | 0.508 | 0.707 | 0.774 |
| PCP_AL | Composition of residues having aliphatic side chains | 0.363 | 0.310 | 0.415 | 0.674 | 0.745 |
| PCP_CY | Composition of residues having cyclic side chains | 0.053 | 0.058 | 0.048 | 0.559 | 0.578 |
| PCP_AR | Composition of aromatic residues | 0.092 | 0.092 | 0.093 | 0.479 | 0.490 |
| PCP_AC | Composition of acidic residues | 0.087 | 0.088 | 0.087 | 0.519 | 0.513 |
| PCP_BS | Composition of basic residues | 0.151 | 0.156 | 0.146 | 0.532 | 0.554 |
| PCP_NE_ph | Composition of neutral residues based on pH | 0.762 | 0.756 | 0.767 | 0.531 | 0.541 |
| PCP_HB | Composition of hydrophobic residues | 0.505 | 0.494 | 0.515 | 0.546 | 0.559 |
| PCP_HL | Composition of hydrophilic residues | 0.250 | 0.262 | 0.237 | 0.570 | 0.594 |
| PCP_NT | Composition of neutral residues | 0.308 | 0.304 | 0.311 | 0.505 | 0.509 |
| PCP_HX | Composition of hydroxylic residues | 0.108 | 0.102 | 0.115 | 0.536 | 0.558 |
| PCP_SC | Composition of residues having sulfur content | 0.125 | 0.183 | 0.067 | 0.840 | 0.881 |
| PCP_SS_HE | Composition of residues in secondary structure (Helix) | 0.373 | 0.321 | 0.424 | 0.693 | 0.748 |
| PCP_SS_ST | Composition of residues in secondary structure (Strands) | 0.335 | 0.369 | 0.301 | 0.654 | 0.697 |
| PCP_SS_CO | Composition of residues in secondary structure (Coil) | 0.292 | 0.309 | 0.274 | 0.591 | 0.620 |
| PCP_SA_BU | Composition of residues in solvent accessibility (Buried) | 0.468 | 0.472 | 0.463 | 0.523 | 0.524 |
| PCP_SA_EX | Composition of residues in solvent accessibility (Exposed) | 0.292 | 0.300 | 0.283 | 0.544 | 0.554 |
| PCP_SA_IN | Composition of residues in solvent accessibility (Intermediate) | 0.241 | 0.236 | 0.246 | 0.520 | 0.526 |
| PCP_TN | Composition of tiny residues | 0.310 | 0.369 | 0.251 | 0.731 | 0.795 |
| PCP_SM | Composition of small residues | 0.548 | 0.602 | 0.494 | 0.695 | 0.744 |
| PCP_LR | Composition of large residues | 0.452 | 0.398 | 0.506 | 0.695 | 0.744 |
| PCP_Z1 | Composition of residues having Z1 advanced physico-chemical properties | 0.281 | 0.575 | -0.014 | 0.689 | 0.736 |
| PCP_Z2 | Composition of residues having Z2 advanced physico-chemical properties | -0.493 | -0.482 | -0.504 | 0.494 | 0.513 |
| PCP_Z3 | Composition of residues having Z3 advanced physico-chemical properties | 0.101 | 0.441 | -0.238 | 0.809 | 0.864 |
| PCP_Z4 | Composition of residues having Z4 advanced physico-chemical properties | -0.203 | -0.124 | -0.282 | 0.617 | 0.643 |
| PCP_Z5 | Composition of residues having Z5 advanced physico-chemical properties | -0.165 | -0.377 | 0.046 | 0.830 | 0.879 |

**Supplementary Table S5.2: Physicochemical Property-based Prediction-based prediction of neurotoxic and non-toxic sequences in protein dataset.**

| Feature | Description | Best Threshold | Mean (Neurotoxic) | Mean (Non-toxic) | Accuracy | AUC |
| --- | --- | --- | --- | --- | --- | --- |
| PCP_PC | Composition of positively charged residues | 0.147 | 0.146 | 0.147 | 0.494 | 0.484 |
| PCP_NC | Composition of negatively charged residues | 0.116 | 0.119 | 0.113 | 0.536 | 0.539 |
| PCP_NE | Composition of neutral charged residues | 0.738 | 0.735 | 0.740 | 0.515 | 0.519 |
| PCP_PO | Composition of polar residues | 0.216 | 0.221 | 0.210 | 0.525 | 0.544 |
| PCP_NP | Composition of non-polar residues | 0.477 | 0.465 | 0.489 | 0.568 | 0.586 |
| PCP_AL | Composition of residues having aliphatic side chains | 0.400 | 0.390 | 0.409 | 0.551 | 0.574 |
| PCP_CY | Composition of residues having cyclic side chains | 0.042 | 0.037 | 0.046 | 0.599 | 0.601 |
| PCP_AR | Composition of aromatic residues | 0.080 | 0.079 | 0.082 | 0.512 | 0.512 |
| PCP_AC | Composition of acidic residues | 0.116 | 0.119 | 0.113 | 0.536 | 0.539 |
| PCP_BS | Composition of basic residues | 0.147 | 0.146 | 0.147 | 0.494 | 0.484 |
| PCP_NE_ph | Composition of neutral residues based on pH | 0.738 | 0.735 | 0.740 | 0.515 | 0.519 |
| PCP_HB | Composition of hydrophobic residues | 0.499 | 0.498 | 0.499 | 0.513 | 0.502 |
| PCP_HL | Composition of hydrophilic residues | 0.233 | 0.232 | 0.235 | 0.488 | 0.495 |
| PCP_NT | Composition of neutral residues | 0.330 | 0.325 | 0.334 | 0.535 | 0.554 |
| PCP_HX | Composition of hydroxylic residues | 0.116 | 0.112 | 0.120 | 0.549 | 0.560 |
| PCP_SC | Composition of residues having sulfur content | 0.063 | 0.078 | 0.048 | 0.632 | 0.683 |
| PCP_SS_HE | Composition of residues in secondary structure (Helix) | 0.437 | 0.423 | 0.452 | 0.594 | 0.618 |
| PCP_SS_ST | Composition of residues in secondary structure (Strands) | 0.297 | 0.313 | 0.281 | 0.610 | 0.658 |
| PCP_SS_CO | Composition of residues in secondary structure (Coil) | 0.266 | 0.265 | 0.267 | 0.493 | 0.505 |
| PCP_SA_BU | Composition of residues in solvent accessibility (Buried) | 0.442 | 0.448 | 0.436 | 0.535 | 0.560 |
| PCP_SA_EX | Composition of residues in solvent accessibility (Exposed) | 0.311 | 0.312 | 0.310 | 0.514 | 0.511 |
| PCP_SA_IN | Composition of residues in solvent accessibility (Intermediate) | 0.236 | 0.228 | 0.244 | 0.586 | 0.592 |
| PCP_TN | Composition of tiny residues | 0.234 | 0.242 | 0.227 | 0.542 | 0.565 |
| PCP_SM | Composition of small residues | 0.487 | 0.494 | 0.480 | 0.539 | 0.556 |
| PCP_LR | Composition of large residues | 0.513 | 0.506 | 0.520 | 0.539 | 0.556 |
| PCP_Z1 | Composition of residues having Z1 advanced physico-chemical properties | 0.117 | 0.149 | 0.084 | 0.526 | 0.523 |
| PCP_Z2 | Composition of residues having Z2 advanced physico-chemical properties | -0.482 | -0.510 | -0.454 | 0.550 | 0.566 |
| PCP_Z3 | Composition of residues having Z3 advanced physico-chemical properties | -0.266 | -0.215 | -0.316 | 0.568 | 0.605 |
| PCP_Z4 | Composition of residues having Z4 advanced physico-chemical properties | -0.384 | -0.397 | -0.372 | 0.525 | 0.532 |
| PCP_Z5 | Composition of residues having Z5 advanced physico-chemical properties | 0.047 | -0.022 | 0.116 | 0.671 | 0.753 |

**Supplementary Table S5.3: Physicochemical Property-based Prediction-based prediction of neurotoxic and non-toxic sequences in combined dataset.**

| Feature | Description | Best Threshold | Mean (Neurotoxic) | Mean (Non-toxic) | Accuracy | AUC |
| --- | --- | --- | --- | --- | --- | --- |
| PCP_PC | Composition of positively charged residues | 0.149 | 0.151 | 0.146 | 0.520 | 0.537 |
| PCP_NC | Composition of negatively charged residues | 0.101 | 0.102 | 0.099 | 0.525 | 0.523 |
| PCP_NE | Composition of neutral charged residues | 0.751 | 0.746 | 0.755 | 0.527 | 0.532 |
| PCP_PO | Composition of polar residues | 0.246 | 0.279 | 0.213 | 0.652 | 0.692 |
| PCP_NP | Composition of non-polar residues | 0.459 | 0.419 | 0.499 | 0.632 | 0.696 |
| PCP_AL | Composition of residues having aliphatic side chains | 0.380 | 0.347 | 0.412 | 0.612 | 0.673 |
| PCP_CY | Composition of residues having cyclic side chains | 0.048 | 0.048 | 0.047 | 0.486 | 0.510 |
| PCP_AR | Composition of aromatic residues | 0.087 | 0.086 | 0.088 | 0.500 | 0.497 |
| PCP_AC | Composition of acidic residues | 0.101 | 0.102 | 0.099 | 0.525 | 0.523 |
| PCP_BS | Composition of basic residues | 0.149 | 0.151 | 0.146 | 0.520 | 0.537 |
| PCP_NE_ph | Composition of neutral residues based on pH | 0.751 | 0.746 | 0.755 | 0.527 | 0.532 |
| PCP_HB | Composition of hydrophobic residues | 0.502 | 0.496 | 0.508 | 0.528 | 0.535 |
| PCP_HL | Composition of hydrophilic residues | 0.242 | 0.248 | 0.236 | 0.538 | 0.558 |
| PCP_NT | Composition of neutral residues | 0.318 | 0.314 | 0.322 | 0.509 | 0.527 |
| PCP_HX | Composition of hydroxylic residues | 0.112 | 0.106 | 0.117 | 0.546 | 0.557 |
| PCP_SC | Composition of residues having sulfur content | 0.096 | 0.134 | 0.058 | 0.748 | 0.777 |
| PCP_SS_HE | Composition of residues in secondary structure (Helix) | 0.403 | 0.369 | 0.437 | 0.624 | 0.684 |
| PCP_SS_ST | Composition of residues in secondary structure (Strands) | 0.317 | 0.343 | 0.292 | 0.631 | 0.675 |
| PCP_SS_CO | Composition of residues in secondary structure (Coil) | 0.280 | 0.289 | 0.271 | 0.539 | 0.572 |
| PCP_SA_BU | Composition of residues in solvent accessibility (Buried) | 0.456 | 0.461 | 0.450 | 0.528 | 0.542 |
| PCP_SA_EX | Composition of residues in solvent accessibility (Exposed) | 0.301 | 0.306 | 0.296 | 0.521 | 0.534 |
| PCP_SA_IN | Composition of residues in solvent accessibility (Intermediate) | 0.239 | 0.232 | 0.245 | 0.548 | 0.551 |
| PCP_TN | Composition of tiny residues | 0.274 | 0.309 | 0.240 | 0.654 | 0.690 |
| PCP_SM | Composition of small residues | 0.520 | 0.551 | 0.488 | 0.610 | 0.660 |
| PCP_LR | Composition of large residues | 0.480 | 0.449 | 0.512 | 0.610 | 0.660 |
| PCP_Z1 | Composition of residues having Z1 advanced physico-chemical properties | 0.204 | 0.375 | 0.032 | 0.597 | 0.651 |
| PCP_Z2 | Composition of residues having Z2 advanced physico-chemical properties | -0.488 | -0.495 | -0.480 | 0.526 | 0.521 |
| PCP_Z3 | Composition of residues having Z3 advanced physico-chemical properties | -0.071 | 0.133 | -0.275 | 0.697 | 0.740 |
| PCP_Z4 | Composition of residues having Z4 advanced physico-chemical properties | -0.288 | -0.252 | -0.324 | 0.558 | 0.568 |
| PCP_Z5 | Composition of residues having Z5 advanced physico-chemical properties | -0.066 | -0.210 | 0.079 | 0.722 | 0.800 |

**Note:** The Z4 and Z5 features refer to two of the five “z‐scale” descriptors that summarize amino acid physicochemical properties via a principal component analysis. In many studies, researchers compute five orthogonal (independent) descriptors—commonly labeled z1 through z5, from a large set of individual properties (such as hydrophobicity, steric bulk, polarity/charge ratio, and electronic characteristics). While z1 to z3 typically capture the most dominant trends (for example, hydrophobicity, size, and polarity/charge ratio), z4 and z5 encapsulate more subtle aspects of amino acid chemistry (often related to electronic effects and other nuanced attributes) that can influence peptide behavior. In prediction models, the overall (often averaged) z4 and z5 values of a peptide are used as features that provide an “advanced” characterization of its physico-chemical makeup.

Z1: Lipophilicity

Z2: Steric properties (Steric bulk/Polarizability)

Z3: Electronic properties (Polarity / Charge ratio)

Z4 and Z5: They relate electronegativity, heat of formation, electrophilicity and hardness.

Supplementary Table S6: Performance evaluation of machine learning models on the different datasets using binary profile features.

**Supplementary Table S6.1: Performance evaluation of machine learning models on the peptide dataset using binary profile features.**

| **Classifier** | **Main Dataset** | | | | | | | | | | | |
| --- | --- | --- | --- | --- | --- | --- | --- | --- | --- | --- | --- | --- |
|  | **Coss-validation Dataset** | | | | | | **Independent Dataset** | | | | | |
|  | **SENS** | **SPEC** | **PREC** | **ACC** | **MCC** | **AUC** | **SENS** | **SPEC** | **PREC** | **ACC** | **MCC** | **AUC** |
| **RF** | 0.909 | 0.864 | 0.870 | 0.887 | 0.774 | 0.945 | 0.931 | 0.841 | 0.853 | 0.886 | 0.775 | 0.946 |
| **GB** | 0.870 | 0.863 | 0.864 | 0.867 | 0.733 | 0.921 | 0.851 | 0.847 | 0.847 | 0.849 | 0.698 | 0.922 |
| **ElasticNet** | 0.856 | 0.860 | 0.860 | 0.858 | 0.716 | 0.911 | 0.846 | 0.818 | 0.822 | 0.832 | 0.664 | 0.913 |
| **Lasso regularization** | 0.848 | 0.856 | 0.855 | 0.852 | 0.704 | 0.912 | 0.846 | 0.858 | 0.855 | 0.852 | 0.704 | 0.918 |
| **Ridge regularization** | 0.860 | 0.846 | 0.848 | 0.853 | 0.706 | 0.896 | 0.846 | 0.818 | 0.822 | 0.832 | 0.664 | 0.908 |
| **LR** | 0.843 | 0.806 | 0.813 | 0.825 | 0.650 | 0.896 | 0.869 | 0.847 | 0.849 | 0.858 | 0.715 | 0.909 |
| **SVC (linear)** | 0.838 | 0.787 | 0.798 | 0.813 | 0.626 | 0.878 | 0.834 | 0.824 | 0.825 | 0.829 | 0.658 | 0.890 |
| **SVC (rbf)** | 0.865 | 0.887 | 0.885 | 0.876 | 0.752 | 0.932 | 0.891 | 0.852 | 0.857 | 0.872 | 0.744 | 0.938 |
| **SVC (poly)** | 0.809 | 0.930 | 0.921 | 0.870 | 0.745 | 0.948 | 0.806 | 0.920 | 0.910 | 0.863 | 0.731 | 0.957 |
| **SVC (sigmoid)** | 0.843 | 0.849 | 0.848 | 0.846 | 0.692 | 0.899 | 0.829 | 0.801 | 0.806 | 0.815 | 0.630 | 0.897 |
| **GNB** | 0.818 | 0.785 | 0.792 | 0.801 | 0.603 | 0.859 | 0.771 | 0.795 | 0.789 | 0.783 | 0.567 | 0.830 |
| **DT** | 0.819 | 0.813 | 0.814 | 0.816 | 0.632 | 0.816 | 0.783 | 0.784 | 0.783 | 0.783 | 0.567 | 0.783 |
| **MLP** | 0.853 | 0.813 | 0.821 | 0.833 | 0.667 | 0.907 | 0.886 | 0.852 | 0.856 | 0.869 | 0.738 | 0.925 |
| **AdB** | 0.835 | 0.819 | 0.822 | 0.827 | 0.654 | 0.896 | 0.823 | 0.807 | 0.809 | 0.815 | 0.630 | 0.902 |
| **XGB** | 0.859 | 0.870 | 0.869 | 0.865 | 0.729 | 0.920 | 0.846 | 0.841 | 0.841 | 0.843 | 0.687 | 0.927 |
| **ET** | 0.905 | 0.860 | 0.866 | 0.882 | 0.766 | 0.942 | 0.914 | 0.847 | 0.856 | 0.880 | 0.763 | 0.949 |
| **GNB** | 0.926 | 0.829 | 0.844 | 0.877 | 0.758 | 0.947 | 0.926 | 0.807 | 0.827 | 0.866 | 0.738 | 0.946 |

**Supplementary Table S6.2: Performance evaluation of machine learning models on the protein dataset using binary profile features.**

| **Classifier** | Protein Dataset | | | | | | | | | | | |
| --- | --- | --- | --- | --- | --- | --- | --- | --- | --- | --- | --- | --- |
|  | Coss-validation Dataset | | | | | | Independent Dataset | | | | | |
|  | **SENS** | **SPEC** | **PREC** | **ACC** | **MCC** | **AUC** | **SENS** | **SPEC** | **PREC** | **ACC** | **MCC** | **AUC** |
| **RF** | 0.690 | 0.668 | 0.675 | 0.679 | 0.358 | 0.741 | 0.703 | 0.697 | 0.699 | 0.700 | 0.400 | 0.768 |
| **GB** | 0.655 | 0.719 | 0.700 | 0.687 | 0.375 | 0.736 | 0.632 | 0.723 | 0.695 | 0.677 | 0.356 | 0.729 |
| **ElasticNet** | 0.642 | 0.731 | 0.704 | 0.686 | 0.374 | 0.749 | 0.639 | 0.735 | 0.707 | 0.687 | 0.376 | 0.751 |
| **Lasso regularization** | 0.661 | 0.703 | 0.690 | 0.682 | 0.365 | 0.743 | 0.658 | 0.755 | 0.729 | 0.706 | 0.415 | 0.756 |
| **Ridge regularization** | 0.666 | 0.726 | 0.708 | 0.696 | 0.393 | 0.726 | 0.658 | 0.723 | 0.703 | 0.690 | 0.381 | 0.756 |
| **LR** | 0.666 | 0.666 | 0.666 | 0.666 | 0.332 | 0.730 | 0.658 | 0.697 | 0.685 | 0.677 | 0.355 | 0.734 |
| **SVC (linear)** | 0.621 | 0.676 | 0.657 | 0.648 | 0.297 | 0.707 | 0.619 | 0.697 | 0.671 | 0.658 | 0.317 | 0.706 |
| **SVC (rbf)** | 0.663 | 0.747 | 0.724 | 0.705 | 0.411 | 0.757 | 0.658 | 0.742 | 0.718 | 0.700 | 0.401 | 0.761 |
| **SVC (poly)** | 0.661 | 0.753 | 0.728 | 0.707 | 0.416 | 0.761 | 0.639 | 0.742 | 0.712 | 0.690 | 0.383 | 0.750 |
| **SVC (sigmoid)** | 0.650 | 0.697 | 0.682 | 0.673 | 0.347 | 0.738 | 0.658 | 0.710 | 0.694 | 0.684 | 0.368 | 0.752 |
| **GNB** | 0.352 | 0.852 | 0.703 | 0.602 | 0.235 | 0.676 | 0.503 | 0.806 | 0.722 | 0.655 | 0.325 | 0.743 |
| **DT** | 0.595 | 0.577 | 0.585 | 0.586 | 0.173 | 0.586 | 0.581 | 0.555 | 0.566 | 0.568 | 0.136 | 0.568 |
| **MLP** | 0.652 | 0.687 | 0.676 | 0.669 | 0.339 | 0.727 | 0.632 | 0.690 | 0.671 | 0.661 | 0.323 | 0.727 |
| **AdB** | 0.652 | 0.663 | 0.659 | 0.657 | 0.315 | 0.725 | 0.639 | 0.735 | 0.707 | 0.687 | 0.376 | 0.749 |
| **GNB** | 0.784 | 0.555 | 0.638 | 0.669 | 0.348 | 0.745 | 0.806 | 0.548 | 0.641 | 0.677 | 0.367 | 0.731 |
| **XGB** | 0.639 | 0.727 | 0.701 | 0.683 | 0.368 | 0.734 | 0.600 | 0.761 | 0.715 | 0.681 | 0.366 | 0.738 |
| **ET** | 0.681 | 0.703 | 0.696 | 0.692 | 0.384 | 0.738 | 0.645 | 0.716 | 0.694 | 0.681 | 0.362 | 0.750 |

**Supplementary Table S6.3: Performance evaluation of machine learning models on the combined dataset using binary profile features.**

| **Classifier** | **Combined Dataset** | | | | | | | | | | | |
| --- | --- | --- | --- | --- | --- | --- | --- | --- | --- | --- | --- | --- |
|  | **Coss-validation Dataset** | | | | | | **Independent Dataset** | | | | | |
|  | **SENS** | **SPEC** | **PREC** | **ACC** | **MCC** | **AUC** | **SENS** | **SPEC** | **PREC** | **ACC** | **MCC** | **AUC** |
| **RF** | 0.830 | 0.692 | 0.729 | 0.761 | 0.527 | 0.830 | 0.833 | 0.683 | 0.724 | 0.758 | 0.522 | 0.841 |
| **GB** | 0.822 | 0.668 | 0.712 | 0.745 | 0.496 | 0.805 | 0.824 | 0.650 | 0.701 | 0.737 | 0.481 | 0.800 |
| **ElasticNet** | 0.789 | 0.678 | 0.710 | 0.733 | 0.469 | 0.796 | 0.794 | 0.671 | 0.706 | 0.732 | 0.468 | 0.804 |
| **Lasso regularization** | 0.782 | 0.684 | 0.713 | 0.733 | 0.469 | 0.798 | 0.788 | 0.686 | 0.714 | 0.737 | 0.476 | 0.810 |
| **Ridge regularization** | 0.756 | 0.689 | 0.709 | 0.723 | 0.446 | 0.789 | 0.764 | 0.710 | 0.724 | 0.737 | 0.474 | 0.809 |
| **LR** | 0.731 | 0.696 | 0.706 | 0.713 | 0.427 | 0.783 | 0.745 | 0.713 | 0.721 | 0.729 | 0.459 | 0.807 |
| **SVC (linear)** | 0.738 | 0.695 | 0.708 | 0.716 | 0.433 | 0.782 | 0.764 | 0.692 | 0.712 | 0.728 | 0.457 | 0.797 |
| **SVC (rbf)** | 0.834 | 0.671 | 0.717 | 0.752 | 0.511 | 0.819 | 0.848 | 0.662 | 0.714 | 0.755 | 0.519 | 0.827 |
| **SVC (poly)** | 0.859 | 0.675 | 0.726 | 0.767 | 0.543 | 0.833 | 0.867 | 0.656 | 0.715 | 0.761 | 0.534 | 0.831 |
| **SVC (sigmoid)** | 0.775 | 0.660 | 0.695 | 0.717 | 0.438 | 0.785 | 0.797 | 0.653 | 0.696 | 0.725 | 0.454 | 0.799 |
| **SGDC** | 0.638 | 0.715 | 0.692 | 0.677 | 0.355 | 0.722 | 0.518 | 0.785 | 0.707 | 0.652 | 0.315 | 0.739 |
| **GNB** | 0.812 | 0.579 | 0.659 | 0.696 | 0.403 | 0.755 | 0.861 | 0.601 | 0.683 | 0.731 | 0.478 | 0.795 |
| **DT** | 0.675 | 0.670 | 0.672 | 0.672 | 0.345 | 0.672 | 0.664 | 0.662 | 0.662 | 0.663 | 0.325 | 0.664 |
| **MLP** | 0.725 | 0.717 | 0.719 | 0.721 | 0.442 | 0.797 | 0.727 | 0.767 | 0.757 | 0.747 | 0.495 | 0.820 |
| **AdB** | 0.761 | 0.671 | 0.699 | 0.716 | 0.434 | 0.783 | 0.776 | 0.686 | 0.711 | 0.731 | 0.463 | 0.796 |
| **GNB** | 0.849 | 0.680 | 0.726 | 0.764 | 0.536 | 0.816 | 0.833 | 0.662 | 0.711 | 0.747 | 0.502 | 0.808 |
| **XGB** | 0.819 | 0.665 | 0.710 | 0.742 | 0.490 | 0.804 | 0.842 | 0.656 | 0.709 | 0.749 | 0.507 | 0.812 |
| **ET** | 0.804 | 0.727 | 0.747 | 0.766 | 0.533 | 0.836 | 0.797 | 0.731 | 0.747 | 0.764 | 0.529 | 0.846 |

Supplementary Table S7: Performance evaluation of machine learning models on the protein datasets using PSSM profiles.

**Supplementary Table S7.1: Performance evaluation of machine learning models on the protein dataset using *aac_pssm* profile features.**

| **Classifier** | **Coss-validation Dataset** | | | | | | **Independent Dataset** | | | | | |
| --- | --- | --- | --- | --- | --- | --- | --- | --- | --- | --- | --- | --- |
|  | **SENS** | **SPEC** | **PREC** | **ACC** | **MCC** | **AUC** | **SENS** | **SPEC** | **PREC** | **ACC** | **MCC** | **AUC** |
| **RF** | 0.789 | 0.751 | 0.760 | 0.770 | 0.540 | 0.837 | 0.787 | 0.755 | 0.763 | 0.771 | 0.542 | 0.830 |
| **GB** | 0.765 | 0.752 | 0.756 | 0.758 | 0.517 | 0.827 | 0.768 | 0.781 | 0.778 | 0.774 | 0.548 | 0.830 |
| **ElasticNet** | 0.734 | 0.709 | 0.717 | 0.721 | 0.443 | 0.778 | 0.800 | 0.710 | 0.734 | 0.755 | 0.512 | 0.808 |
| **Lasso regularization** | 0.760 | 0.717 | 0.729 | 0.738 | 0.477 | 0.797 | 0.781 | 0.774 | 0.776 | 0.777 | 0.555 | 0.838 |
| **Ridge regularization** | 0.761 | 0.714 | 0.727 | 0.737 | 0.475 | 0.795 | 0.781 | 0.781 | 0.781 | 0.781 | 0.561 | 0.838 |
| **LR** | 0.765 | 0.723 | 0.735 | 0.744 | 0.488 | 0.800 | 0.781 | 0.755 | 0.761 | 0.768 | 0.536 | 0.839 |
| **SVC (linear)** | 0.769 | 0.725 | 0.737 | 0.747 | 0.495 | 0.801 | 0.768 | 0.755 | 0.758 | 0.761 | 0.523 | 0.837 |
| **SVC (rbf)** | 0.785 | 0.757 | 0.765 | 0.771 | 0.543 | 0.835 | 0.781 | 0.742 | 0.752 | 0.761 | 0.523 | 0.841 |
| **SVC (poly)** | 0.802 | 0.720 | 0.742 | 0.761 | 0.523 | 0.819 | 0.787 | 0.742 | 0.753 | 0.765 | 0.530 | 0.839 |
| **SVC (sigmoid)** | 0.440 | 0.417 | 0.431 | 0.429 | -0.142 | 0.605 | 0.419 | 0.335 | 0.387 | 0.377 | -0.246 | 0.665 |
| **SGDC** | 0.577 | 0.691 | 0.652 | 0.634 | 0.270 | 0.642 | 0.497 | 0.897 | 0.828 | 0.697 | 0.429 | 0.727 |
| **GNB** | 0.739 | 0.545 | 0.620 | 0.642 | 0.290 | 0.710 | 0.794 | 0.574 | 0.651 | 0.684 | 0.377 | 0.744 |
| **DT** | 0.673 | 0.720 | 0.707 | 0.696 | 0.393 | 0.696 | 0.710 | 0.665 | 0.679 | 0.687 | 0.375 | 0.687 |
| **MLP** | 0.769 | 0.773 | 0.773 | 0.771 | 0.543 | 0.838 | 0.787 | 0.781 | 0.782 | 0.784 | 0.568 | 0.848 |
| **AdB** | 0.748 | 0.736 | 0.740 | 0.742 | 0.485 | 0.804 | 0.742 | 0.735 | 0.737 | 0.739 | 0.477 | 0.804 |
| **GNB** | 0.766 | 0.777 | 0.775 | 0.771 | 0.543 | 0.840 | 0.774 | 0.794 | 0.789 | 0.784 | 0.568 | 0.847 |
| **XGB** | 0.769 | 0.764 | 0.766 | 0.767 | 0.533 | 0.832 | 0.768 | 0.761 | 0.763 | 0.765 | 0.529 | 0.840 |
| **ET** | 0.761 | 0.759 | 0.760 | 0.760 | 0.520 | 0.825 | 0.761 | 0.761 | 0.761 | 0.761 | 0.523 | 0.833 |

**Supplementary Table S7.2: Performance evaluation of machine learning models on the protein dataset using *aac_pssm +* AAC composition profile features.**

| **Classifier** | **Coss-validation Dataset** | | | | | | **Independent Dataset** | | | | | |
| --- | --- | --- | --- | --- | --- | --- | --- | --- | --- | --- | --- | --- |
|  | **SENS** | **SPEC** | **PREC** | **ACC** | **MCC** | **AUC** | **SENS** | **SPEC** | **PREC** | **ACC** | **MCC** | **AUC** |
| **RF** | 0.762 | 0.821 | 0.809 | 0.792 | 0.584 | 0.857 | 0.768 | 0.800 | 0.793 | 0.784 | 0.568 | 0.855 |
| **GB** | 0.778 | 0.802 | 0.796 | 0.790 | 0.580 | 0.853 | 0.768 | 0.781 | 0.778 | 0.774 | 0.548 | 0.841 |
| **ElasticNet** | 0.783 | 0.768 | 0.771 | 0.775 | 0.551 | 0.825 | 0.748 | 0.729 | 0.734 | 0.739 | 0.478 | 0.813 |
| **Lasso regularization** | 0.778 | 0.768 | 0.770 | 0.773 | 0.546 | 0.827 | 0.748 | 0.742 | 0.744 | 0.745 | 0.490 | 0.814 |
| **Ridge regularization** | 0.786 | 0.758 | 0.764 | 0.772 | 0.545 | 0.828 | 0.748 | 0.755 | 0.753 | 0.752 | 0.503 | 0.814 |
| **LR** | 0.780 | 0.760 | 0.764 | 0.770 | 0.540 | 0.824 | 0.742 | 0.710 | 0.719 | 0.726 | 0.452 | 0.808 |
| **SVC (linear)** | 0.765 | 0.771 | 0.769 | 0.768 | 0.536 | 0.827 | 0.755 | 0.748 | 0.750 | 0.752 | 0.503 | 0.817 |
| **SVC (rbf)** | 0.814 | 0.802 | 0.804 | 0.808 | 0.616 | 0.871 | 0.800 | 0.787 | 0.790 | 0.794 | 0.587 | 0.861 |
| **SVC (poly)** | 0.811 | 0.794 | 0.797 | 0.802 | 0.604 | 0.864 | 0.794 | 0.781 | 0.783 | 0.787 | 0.574 | 0.849 |
| **SVC (sigmoid)** | 0.757 | 0.710 | 0.722 | 0.733 | 0.467 | 0.779 | 0.735 | 0.697 | 0.708 | 0.716 | 0.433 | 0.752 |
| **SGDC** | 0.524 | 0.805 | 0.728 | 0.665 | 0.343 | 0.666 | 0.303 | 0.968 | 0.904 | 0.635 | 0.363 | 0.635 |
| **GNB** | 0.772 | 0.650 | 0.687 | 0.711 | 0.425 | 0.789 | 0.794 | 0.594 | 0.661 | 0.694 | 0.395 | 0.777 |
| **DT** | 0.683 | 0.702 | 0.695 | 0.692 | 0.385 | 0.692 | 0.619 | 0.690 | 0.667 | 0.655 | 0.310 | 0.655 |
| **MLP** | 0.793 | 0.774 | 0.778 | 0.784 | 0.567 | 0.858 | 0.774 | 0.787 | 0.784 | 0.781 | 0.561 | 0.859 |
| **AdB** | 0.748 | 0.740 | 0.742 | 0.744 | 0.488 | 0.817 | 0.716 | 0.735 | 0.730 | 0.726 | 0.452 | 0.793 |
| **GNB** | 0.790 | 0.618 | 0.673 | 0.704 | 0.413 | 0.666 | 0.826 | 0.600 | 0.674 | 0.713 | 0.437 | 0.721 |
| **XGB** | 0.778 | 0.782 | 0.781 | 0.780 | 0.561 | 0.854 | 0.755 | 0.774 | 0.770 | 0.765 | 0.529 | 0.839 |
| **ET** | 0.799 | 0.808 | 0.806 | 0.804 | 0.607 | 0.862 | 0.826 | 0.748 | 0.766 | 0.787 | 0.576 | 0.852 |

**Supplementary Table S7.3: Performance evaluation of machine learning models on the protein dataset using *mepd_pssm* composition profile features.**

| **Classifier** | **Coss-validation Dataset** | | | | | | **Independent Dataset** | | | | | |
| --- | --- | --- | --- | --- | --- | --- | --- | --- | --- | --- | --- | --- |
|  | **SENS** | **SPEC** | **PREC** | **ACC** | **MCC** | **AUC** | **SENS** | **SPEC** | **PREC** | **ACC** | **MCC** | **AUC** |
| **RF** | 0.789 | 0.751 | 0.760 | 0.770 | 0.540 | 0.837 | 0.787 | 0.755 | 0.763 | 0.771 | 0.542 | 0.830 |
| **GB** | 0.765 | 0.752 | 0.756 | 0.758 | 0.517 | 0.827 | 0.768 | 0.781 | 0.778 | 0.774 | 0.548 | 0.830 |
| **ElasticNet** | 0.734 | 0.709 | 0.717 | 0.721 | 0.443 | 0.778 | 0.800 | 0.710 | 0.734 | 0.755 | 0.512 | 0.808 |
| **Lasso regularization** | 0.760 | 0.717 | 0.729 | 0.738 | 0.477 | 0.797 | 0.781 | 0.774 | 0.776 | 0.777 | 0.555 | 0.838 |
| **Ridge regularization** | 0.761 | 0.714 | 0.727 | 0.737 | 0.475 | 0.795 | 0.781 | 0.781 | 0.781 | 0.781 | 0.561 | 0.838 |
| **LR** | 0.765 | 0.723 | 0.735 | 0.744 | 0.488 | 0.800 | 0.781 | 0.755 | 0.761 | 0.768 | 0.536 | 0.839 |
| **SVC (linear)** | 0.769 | 0.725 | 0.737 | 0.747 | 0.495 | 0.801 | 0.768 | 0.755 | 0.758 | 0.761 | 0.523 | 0.837 |
| **SVC (rbf)** | 0.785 | 0.757 | 0.765 | 0.771 | 0.543 | 0.835 | 0.781 | 0.742 | 0.752 | 0.761 | 0.523 | 0.841 |
| **SVC (poly)** | 0.802 | 0.720 | 0.742 | 0.761 | 0.523 | 0.819 | 0.787 | 0.742 | 0.753 | 0.765 | 0.530 | 0.839 |
| **SVC (sigmoid)** | 0.440 | 0.417 | 0.431 | 0.429 | -0.142 | 0.605 | 0.419 | 0.335 | 0.387 | 0.377 | -0.246 | 0.665 |
| **SGDC** | 0.577 | 0.691 | 0.652 | 0.634 | 0.270 | 0.642 | 0.497 | 0.897 | 0.828 | 0.697 | 0.429 | 0.727 |
| **GNB** | 0.739 | 0.545 | 0.620 | 0.642 | 0.290 | 0.710 | 0.794 | 0.574 | 0.651 | 0.684 | 0.377 | 0.744 |
| **DT** | 0.673 | 0.720 | 0.707 | 0.696 | 0.393 | 0.696 | 0.710 | 0.665 | 0.679 | 0.687 | 0.375 | 0.687 |
| **MLP** | 0.769 | 0.773 | 0.773 | 0.771 | 0.543 | 0.838 | 0.787 | 0.781 | 0.782 | 0.784 | 0.568 | 0.848 |
| **AdB** | 0.748 | 0.736 | 0.740 | 0.742 | 0.485 | 0.804 | 0.742 | 0.735 | 0.737 | 0.739 | 0.477 | 0.804 |
| **GNB** | 0.766 | 0.777 | 0.775 | 0.771 | 0.543 | 0.840 | 0.774 | 0.794 | 0.789 | 0.784 | 0.568 | 0.847 |
| **XGB** | 0.769 | 0.764 | 0.766 | 0.767 | 0.533 | 0.832 | 0.768 | 0.761 | 0.763 | 0.765 | 0.529 | 0.840 |
| **ET** | 0.761 | 0.759 | 0.760 | 0.760 | 0.520 | 0.825 | 0.761 | 0.761 | 0.761 | 0.761 | 0.523 | 0.833 |

**Supplementary Table S7.4: Performance evaluation of machine learning models on the protein dataset using *mepd_pssm +* AAC composition profile features.**

| **Classifier** | **Coss-validation Dataset** | | | | | | **Independent Dataset** | | | | | |
| --- | --- | --- | --- | --- | --- | --- | --- | --- | --- | --- | --- | --- |
|  | **SENS** | **SPEC** | **PREC** | **ACC** | **MCC** | **AUC** | **SENS** | **SPEC** | **PREC** | **ACC** | **MCC** | **AUC** |
| **RF** | 0.762 | 0.821 | 0.809 | 0.792 | 0.584 | 0.857 | 0.768 | 0.800 | 0.793 | 0.784 | 0.568 | 0.855 |
| **GB** | 0.778 | 0.802 | 0.796 | 0.790 | 0.580 | 0.853 | 0.768 | 0.781 | 0.778 | 0.774 | 0.548 | 0.841 |
| **ElasticNet** | 0.783 | 0.768 | 0.771 | 0.775 | 0.551 | 0.825 | 0.748 | 0.729 | 0.734 | 0.739 | 0.478 | 0.813 |
| **Lasso regularization** | 0.778 | 0.768 | 0.770 | 0.773 | 0.546 | 0.827 | 0.748 | 0.742 | 0.744 | 0.745 | 0.490 | 0.814 |
| **Ridge regularization** | 0.786 | 0.758 | 0.764 | 0.772 | 0.545 | 0.828 | 0.748 | 0.755 | 0.753 | 0.752 | 0.503 | 0.814 |
| **LR** | 0.780 | 0.760 | 0.764 | 0.770 | 0.540 | 0.824 | 0.742 | 0.710 | 0.719 | 0.726 | 0.452 | 0.808 |
| **SVC (linear)** | 0.765 | 0.771 | 0.769 | 0.768 | 0.536 | 0.827 | 0.755 | 0.748 | 0.750 | 0.752 | 0.503 | 0.817 |
| **SVC (rbf)** | 0.814 | 0.802 | 0.804 | 0.808 | 0.616 | 0.871 | 0.800 | 0.787 | 0.790 | 0.794 | 0.587 | 0.861 |
| **SVC (poly)** | 0.811 | 0.794 | 0.797 | 0.802 | 0.604 | 0.864 | 0.794 | 0.781 | 0.783 | 0.787 | 0.574 | 0.849 |
| **SVC (sigmoid)** | 0.757 | 0.710 | 0.722 | 0.733 | 0.467 | 0.779 | 0.735 | 0.697 | 0.708 | 0.716 | 0.433 | 0.752 |
| **SGDC** | 0.524 | 0.805 | 0.728 | 0.665 | 0.343 | 0.666 | 0.303 | 0.968 | 0.904 | 0.635 | 0.363 | 0.635 |
| **GNB** | 0.772 | 0.650 | 0.687 | 0.711 | 0.425 | 0.789 | 0.794 | 0.594 | 0.661 | 0.694 | 0.395 | 0.777 |
| **DT** | 0.683 | 0.702 | 0.695 | 0.692 | 0.385 | 0.692 | 0.619 | 0.690 | 0.667 | 0.655 | 0.310 | 0.655 |
| **MLP** | 0.793 | 0.774 | 0.778 | 0.784 | 0.567 | 0.858 | 0.774 | 0.787 | 0.784 | 0.781 | 0.561 | 0.859 |
| **AdB** | 0.748 | 0.740 | 0.742 | 0.744 | 0.488 | 0.817 | 0.716 | 0.735 | 0.730 | 0.726 | 0.452 | 0.793 |
| **GNB** | 0.790 | 0.618 | 0.673 | 0.704 | 0.413 | 0.666 | 0.826 | 0.600 | 0.674 | 0.713 | 0.437 | 0.721 |
| **XGB** | 0.778 | 0.782 | 0.781 | 0.780 | 0.561 | 0.854 | 0.755 | 0.774 | 0.770 | 0.765 | 0.529 | 0.839 |
| **ET** | 0.799 | 0.808 | 0.806 | 0.804 | 0.607 | 0.862 | 0.826 | 0.748 | 0.766 | 0.787 | 0.576 | 0.852 |

**Supplementary Table S7.5: Performance evaluation of machine learning models on the protein dataset using *pssm_composition* composition profile features.**

| **Classifier** | **Coss-validation Dataset** | | | | | | **Independent Dataset** | | | | | |
| --- | --- | --- | --- | --- | --- | --- | --- | --- | --- | --- | --- | --- |
|  | **SENS** | **SPEC** | **PREC** | **ACC** | **MCC** | **AUC** | **SENS** | **SPEC** | **PREC** | **ACC** | **MCC** | **AUC** |
| **RF** | 0.768 | 0.772 | 0.771 | 0.770 | 0.540 | 0.841 | 0.806 | 0.735 | 0.753 | 0.771 | 0.543 | 0.851 |
| **GB** | 0.756 | 0.757 | 0.758 | 0.757 | 0.514 | 0.834 | 0.787 | 0.755 | 0.763 | 0.771 | 0.542 | 0.837 |
| **ElasticNet** | 0.774 | 0.613 | 0.668 | 0.694 | 0.393 | 0.759 | 0.781 | 0.729 | 0.742 | 0.755 | 0.510 | 0.829 |
| **Lasso regularization** | 0.779 | 0.659 | 0.696 | 0.719 | 0.441 | 0.786 | 0.787 | 0.735 | 0.748 | 0.761 | 0.523 | 0.832 |
| **Ridge regularization** | 0.781 | 0.730 | 0.743 | 0.755 | 0.511 | 0.814 | 0.781 | 0.729 | 0.742 | 0.755 | 0.510 | 0.832 |
| **LR** | 0.756 | 0.718 | 0.729 | 0.737 | 0.475 | 0.794 | 0.729 | 0.723 | 0.724 | 0.726 | 0.452 | 0.813 |
| **SVC (linear)** | 0.732 | 0.689 | 0.703 | 0.711 | 0.422 | 0.762 | 0.742 | 0.690 | 0.706 | 0.716 | 0.433 | 0.803 |
| **SVC (rbf)** | 0.802 | 0.748 | 0.761 | 0.775 | 0.550 | 0.841 | 0.832 | 0.729 | 0.754 | 0.781 | 0.564 | 0.842 |
| **SVC (poly)** | 0.806 | 0.675 | 0.713 | 0.741 | 0.485 | 0.796 | 0.794 | 0.690 | 0.719 | 0.742 | 0.486 | 0.799 |
| **SVC (sigmoid)** | 0.800 | 0.073 | 0.464 | 0.437 | -0.185 | 0.619 | 0.839 | 0.045 | 0.468 | 0.442 | -0.191 | 0.684 |
| **GNB** | 0.760 | 0.458 | 0.584 | 0.609 | 0.228 | 0.629 | 0.800 | 0.523 | 0.626 | 0.661 | 0.336 | 0.671 |
| **DT** | 0.645 | 0.670 | 0.662 | 0.658 | 0.315 | 0.658 | 0.632 | 0.710 | 0.685 | 0.671 | 0.343 | 0.671 |
| **MLP** | 0.768 | 0.693 | 0.715 | 0.730 | 0.462 | 0.792 | 0.690 | 0.755 | 0.738 | 0.723 | 0.446 | 0.819 |
| **AdB** | 0.681 | 0.741 | 0.725 | 0.711 | 0.423 | 0.777 | 0.781 | 0.703 | 0.725 | 0.742 | 0.485 | 0.809 |
| **GNB** | 0.632 | 0.748 | 0.715 | 0.690 | 0.382 | 0.595 | 0.619 | 0.787 | 0.744 | 0.703 | 0.412 | 0.621 |
| **XGB** | 0.748 | 0.761 | 0.758 | 0.754 | 0.509 | 0.834 | 0.800 | 0.748 | 0.761 | 0.774 | 0.549 | 0.835 |
| **ET** | 0.787 | 0.778 | 0.781 | 0.783 | 0.565 | 0.839 | 0.800 | 0.748 | 0.761 | 0.774 | 0.549 | 0.842 |

**Supplementary Table S7.6: Performance evaluation of machine learning models on the protein dataset using *pssm_composition +* AAC composition profile features.**

| **Classifier** | **Coss-validation Dataset** | | | | | | **Independent Dataset** | | | | | |
| --- | --- | --- | --- | --- | --- | --- | --- | --- | --- | --- | --- | --- |
|  | **SENS** | **SPEC** | **PREC** | **ACC** | **MCC** | **AUC** | **SENS** | **SPEC** | **PREC** | **ACC** | **MCC** | **AUC** |
| **RF** | 0.717 | 0.695 | 0.701 | 0.706 | 0.412 | 0.770 | 0.671 | 0.632 | 0.646 | 0.652 | 0.303 | 0.700 |
| **GB** | 0.769 | 0.790 | 0.785 | 0.779 | 0.559 | 0.848 | 0.742 | 0.735 | 0.737 | 0.739 | 0.477 | 0.822 |
| **ElasticNet** | 0.767 | 0.769 | 0.768 | 0.768 | 0.536 | 0.827 | 0.755 | 0.735 | 0.741 | 0.745 | 0.490 | 0.811 |
| **Lasso regularization** | 0.778 | 0.771 | 0.772 | 0.775 | 0.549 | 0.828 | 0.748 | 0.729 | 0.734 | 0.739 | 0.478 | 0.815 |
| **Ridge regularization** | 0.752 | 0.776 | 0.770 | 0.764 | 0.528 | 0.818 | 0.774 | 0.690 | 0.714 | 0.732 | 0.466 | 0.800 |
| **LR** | 0.697 | 0.705 | 0.702 | 0.701 | 0.402 | 0.766 | 0.671 | 0.632 | 0.646 | 0.652 | 0.303 | 0.731 |
| **SVC (linear)** | 0.673 | 0.703 | 0.693 | 0.688 | 0.377 | 0.740 | 0.671 | 0.619 | 0.638 | 0.645 | 0.291 | 0.708 |
| **SVC (rbf)** | 0.756 | 0.766 | 0.763 | 0.761 | 0.522 | 0.838 | 0.781 | 0.729 | 0.742 | 0.755 | 0.510 | 0.829 |
| **SVC (poly)** | 0.749 | 0.740 | 0.742 | 0.745 | 0.490 | 0.795 | 0.723 | 0.729 | 0.727 | 0.726 | 0.452 | 0.782 |
| **SVC (sigmoid)** | 0.215 | 0.787 | 0.502 | 0.502 | 0.003 | 0.480 | 0.123 | 0.865 | 0.475 | 0.494 | -0.019 | 0.441 |
| **GNB** | 0.612 | 0.489 | 0.544 | 0.550 | 0.101 | 0.572 | 0.568 | 0.439 | 0.503 | 0.503 | 0.007 | 0.526 |
| **DT** | 0.678 | 0.671 | 0.673 | 0.674 | 0.349 | 0.674 | 0.677 | 0.684 | 0.682 | 0.681 | 0.361 | 0.681 |
| **MLP** | 0.693 | 0.779 | 0.758 | 0.736 | 0.473 | 0.808 | 0.735 | 0.690 | 0.704 | 0.713 | 0.426 | 0.797 |
| **AdB** | 0.725 | 0.745 | 0.739 | 0.735 | 0.470 | 0.795 | 0.716 | 0.677 | 0.689 | 0.697 | 0.394 | 0.771 |
| **GNB** | 0.707 | 0.598 | 0.637 | 0.653 | 0.307 | 0.503 | 0.684 | 0.587 | 0.624 | 0.635 | 0.272 | 0.503 |
| **XGB** | 0.777 | 0.795 | 0.791 | 0.786 | 0.572 | 0.847 | 0.787 | 0.684 | 0.713 | 0.735 | 0.473 | 0.826 |
| **ET** | 0.676 | 0.676 | 0.675 | 0.676 | 0.352 | 0.711 | 0.677 | 0.581 | 0.618 | 0.629 | 0.259 | 0.670 |

Supplementary Table S8: Performance of best machine learning models on neurotoxic datasets using modlAMP features and combination with compositional features.

**Supplementary Table S8: Performance of best machine learning models on neurotoxic datasets using *modlAMP* features and combination with compositional features.**

| **Dataset** | **Features** | **Classifier** | **Coss-validation Dataset** | | | | | | **Independent Dataset** | | | | | |
| --- | --- | --- | --- | --- | --- | --- | --- | --- | --- | --- | --- | --- | --- | --- |
|  |  |  | **SENS** | **SPEC** | **PPV** | **ACC** | **MCC** | **AUC** | **SENS** | **SPEC** | **PPV** | **ACC** | **MCC** | **AUC** |
| **Peptide** | ***modlAMP*** | **ET** | 0.900 | 0.874 | 0.878 | 0.887 | 0.775 | 0.939 | 0.886 | 0.858 | 0.861 | 0.872 | 0.744 | 0.934 |
|  | ***modlAMP*+AAC** | **ET** | 0.897 | 0.884 | 0.886 | 0.891 | 0.782 | 0.941 | 0.897 | 0.875 | 0.877 | 0.886 | 0.772 | 0.945 |
|  | ***modlAMP*+AAC+DPC** | **ET** | 0.913 | 0.879 | 0.883 | 0.896 | 0.792 | 0.949 | 0.903 | 0.852 | 0.859 | 0.877 | 0.756 | 0.955 |
| **Proteins** | ***modlAMP*** | **RF** | 0.748 | 0.813 | 0.800 | 0.781 | 0.773 | 0.854 | 0.684 | 0.839 | 0.809 | 0.761 | 0.741 | 0.845 |
|  | ***modlAMP*+AAC** | **RF** | 0.776 | 0.842 | 0.831 | 0.809 | 0.802 | 0.877 | 0.787 | 0.852 | 0.841 | 0.819 | 0.813 | 0.872 |
|  | ***modlAMP*+AAC+DPC** | **RF** | 0.790 | 0.832 | 0.825 | 0.811 | 0.807 | 0.879 | 0.748 | 0.839 | 0.823 | 0.794 | 0.784 | 0.856 |
| **Combined** | ***modlAMP*** | **RF** | 0.809 | 0.844 | 0.838 | 0.827 | 0.824 | 0.898 | 0.798 | 0.845 | 0.838 | 0.821 | 0.817 | 0.907 |
|  | ***modlAMP*+AAC** | **RF** | 0.824 | 0.859 | 0.854 | 0.842 | 0.839 | 0.907 | 0.816 | 0.873 | 0.865 | 0.844 | 0.840 | 0.915 |
|  | ***modlAMP*+AAC+DPC** | **RF** | 0.834 | 0.859 | 0.855 | 0.846 | 0.844 | 0.911 | 0.831 | 0.864 | 0.859 | 0.847 | 0.845 | 0.919 |

(SENS: Sensitivity; SPEC: Specificity; PPV: Positive Predictive Value; ACC: Accuracy; MCC: Matthews Correlation Coefficient; AUC: Area Under the Receiver Operating Characteristic Curve)
